# A Genome Scale Transcriptional Regulatory Model of the Human Placenta

**DOI:** 10.1101/2022.09.27.509541

**Authors:** Alison Paquette, Kylia Ahuna, Yeon Mi Hwang, Jocelynn Pearl, Hanna Liao, Paul Shannon, Leena Kadam, Samantha Lapehn, Matthew Bucher, Ryan Roper, Cory Funk, James MacDonald, Theo Bammler, Priyanka Baloni, Heather Brockway, W. Alex Mason, Nicole Bush, Kaja Z Lewinn, Catherine J Karr, John Stamatoyannopoulos, Louis J Muglia, Helen Jones, Yoel Sadovsky, Leslie Myatt, Sheela Sathyanarayana, Nathan D. Price, program collaborators for Environmental influences on Child Health Outcomes

## Abstract

Gene regulation is essential to placental function and fetal development. We report a genome-scale transcriptional regulatory network (TRN) of the human placenta built using digital genomic footprinting and transcriptomic data. We integrated 475 transcriptomes and 12 DNase hypersensitivity datasets from placental samples to globally and quantitatively map transcription factor (TF)-target gene interactions. In an independent dataset, the TRN model predicted target gene expression with an out of sample R^2^ value greater than 0.25 for 74% of target genes. We performed siRNA knockdowns of 4 TFs and achieved concordance between the predicted gene targets in our TRN and differences in expression of knockdowns with an accuracy of >0.7 for 3 of the 4 TFs. Our final model contained 113,158 interactions across 391 TFs and 7,712 target genes and is publicly available. We identified six TFs which were significantly enriched as regulators for genes previously associated with preterm birth.

## INTRODUCTION

The placenta is an essential organ with multiple functions.^1^ It regulates gas, nutrient and waste transport, as well as provides immunological defense. It also performs endocrine functions, producing key hormones and modulating the passage of hormonal and paracrine signals between the mother and developing fetus.^2^ Thus, the placenta modulates the *in utero* environment, impacting fetal development, growth, and health.^3^ This complex organ is composed of various types of trophoblast cells, including extravillous trophoblasts, cytotrophoblasts, and the syncytiotrophoblast, which are a large terminally differentiated, multi-nucleated cell type that is the primary site of nutrient uptake and transport and peptide and steroid hormone synthesis, as well as other cell types including stromal cells, fibroblasts, endothelial cells, and various immune cells.^4^ With its many cell types and highly specialized functions, the placenta has a unique transcriptome,^5^ which including numerous genes uniquely expressed only within the placenta. Fetal genomic variation influences placental epigenomic and transcriptomic interactions,^6^ which change across gestation,^78^ and in response to cues from the maternal environment.^9^ The placental transcriptome reflects the underlying physiological functions of the organ, and understanding these changes in normal and pathological placental tissue can provide insight into the molecular mechanisms into pathophysiology, Perturbations to the placental transcriptome are linked to pregnancy pathologies including preterm birth^10,11^ and preeclampsia.^12^ Previous studies of individual genes have revealed several temporally regulated and cell type specific transcription factors (TFs) that play key roles in discrete developmental stages and functions of the placenta. Throughout pregnancy, other transcription factors including *GCM1, GATA2/3* and *CREB* emerge as unique regulators of specific trophoblast subtypes.^13^ These transcription factors regulate key placental processes such as invasion and nutrient transport that may be linked to adverse placental function and prenatal complications. Further, understanding these transcriptional changes in normal and pathological placentas can provide insights into the molecular mechanisms underlying gestational pathophysiology.

Transcriptional regulatory networks (TRNs) map TFs to the target genes they regulate.^**14**,**15**^ This type of network is context-specific; and requires capturing and quantifying the regulatory machinery within specific cellular populations and across developmental stages.^**16**^ TRNs integrate information about chromatin accessibility (captured through DNaseq or ATAC seq) and predicted transcription factor binding, using sequence motifs from a compendium of TFs and RNA sequencing data to produce genome scale networks.^**17**^ A TRN is made up of nodes (TFs and their target genes) and edges that explain which transcription factors play a role in regulating the target genes. These edges are defined by TF binding within regions of open chromatin and established relationships in gene expression between TFs and target genes. TRNs can be organized into units known as “regulons,” which are composed of each individual TF and the target genes they regulate. We have generated such a network previously in the human brain,^**18**^ which revealed (1) TFs regulated by viral abundance associated with Alzheimer’s disease,^**19**^ (2) TFs that drive gene expression changes associated with bipolar disorder and schizophrenia,^**18**^ (3) regulatory functions for SNPs associated with genetic risks associated with these disorders,^**20**^ and (4) TF-target gene relationships underlying gene expression changes related to Huntington’s Disease.^**21**^ This approach has demonstrated that quantifying transcriptional networks has the potential to illuminate the etiology of a wide variety of pathological disorders. We now extend this approach into the placenta.

Other network biology-based approaches have been previously deployed to gain insight into placental function and prenatal disease, such as Weighted gene co-expression analysis (WGCNA), which is a network strategy that identifies co-regulated gene clusters based on correlated expression.^22^ WGCNA was performed on human placental transcriptome data collected at delivery revealed gene clusters that were involved in core placental functions and were reproducible across trimesters (in mice).^23^ An independent WGCNA constructed on a large number of term placenta transcriptomes revealed modules associated with fetal growth restriction.^24^ Combining co-expression analyses with chromatin accessibility data is essential for a deeper understanding of the underlying transcriptional mechanisms of regulation. A placenta TRN constructed using DNase hypersensitivity data derived from first trimester placental data has been used to model the transformation of embryonic stem cells to trophoblast cells; providing key insights into transcription factors which regulate genes with regions of open chromatin in this context.^25^ An accurate model of the placental transcriptome at the end of gestation, enhanced with DNase hypersensitivity data, will thus provide valuable insight into TF-target gene interactions and serve as a tool for placental gene expression analyses at this timepoint.

We characterized placental transcriptional regulation at term by constructing a genome scale TRN of the human placenta, using DNAse hypersensitivity data and curated transcription factor binding motif databases, paired with RNA sequencing data compiled from 475 placental samples. To test the accuracy of our model, we applied a machine learning approach to evaluate the accuracy of the model at predicting gene expression and performed experimental validation of transcriptional regulation of a core group of TFs in isolated villous cytotrophoblasts. Because previous transcriptomics identified fetal sex as a critical biological variable that impacts the placental transcriptome,^26^ and pathology,^27^ we also characterized sex-specific differences in TF-target gene interactions within the placenta. We hypothesized that our network could reveal core TFs that regulated changes in expression of genes whose placental expression was previously associated with preterm birth. We tested this hypothesis by identifying shared regulation of genes whose placental expression was associated with preterm birth within an independent analysis using aggregated microarray data. This approach can be broadly applied to other types placental ‘omics data (e.g. DNA methylation data), providing researchers a different way to interpret lists of candidate genes and identify transcriptional regulators.

## RESULTS

### Reconstruction of a transcriptional regulatory network model for the human placenta

Our placental TRN was constructed using transcriptomic data from 475 placental samples collected in two independent birth cohorts: at Magee Womens Research Institute (MWRI) in Pittsburgh PA and at the University of Tennessee Medical Center in Memphis TN, as a part of the Conditions Affecting Cognitive Development and Learning in Early Childhood (CANDLE) study. DNase hypersensitivity data was generated in 12 term human placentas from Oregon Health & Science University in Portland OR. Complete covariate information for all placental samples used within the model can be found in **Table 1**. The CANDLE cohort was predominantly Black (56%); whereas the samples from MWRI were predominantly White (85%). Overall, the demographics of RNA sequencing samples in this dataset was 46% Black, 52% White, and 2% other Race groups. The mean gestational age of the MWH samples was 39.6 weeks, and the mean gestational age of the CANDLE cohort was 39.2 weeks. 3% of the samples in the CANDLE cohort were born prematurely (<37 weeks), which we elected to retain in the model to capture routine variation in the placental transcriptome related to gestational age. We excluded samples from pregnancies with preeclampsia and multi-fetal gestations in order to capture the placental transcriptome of infants without complicated deliveries that were potentially medically indicated.

**Table 1:**
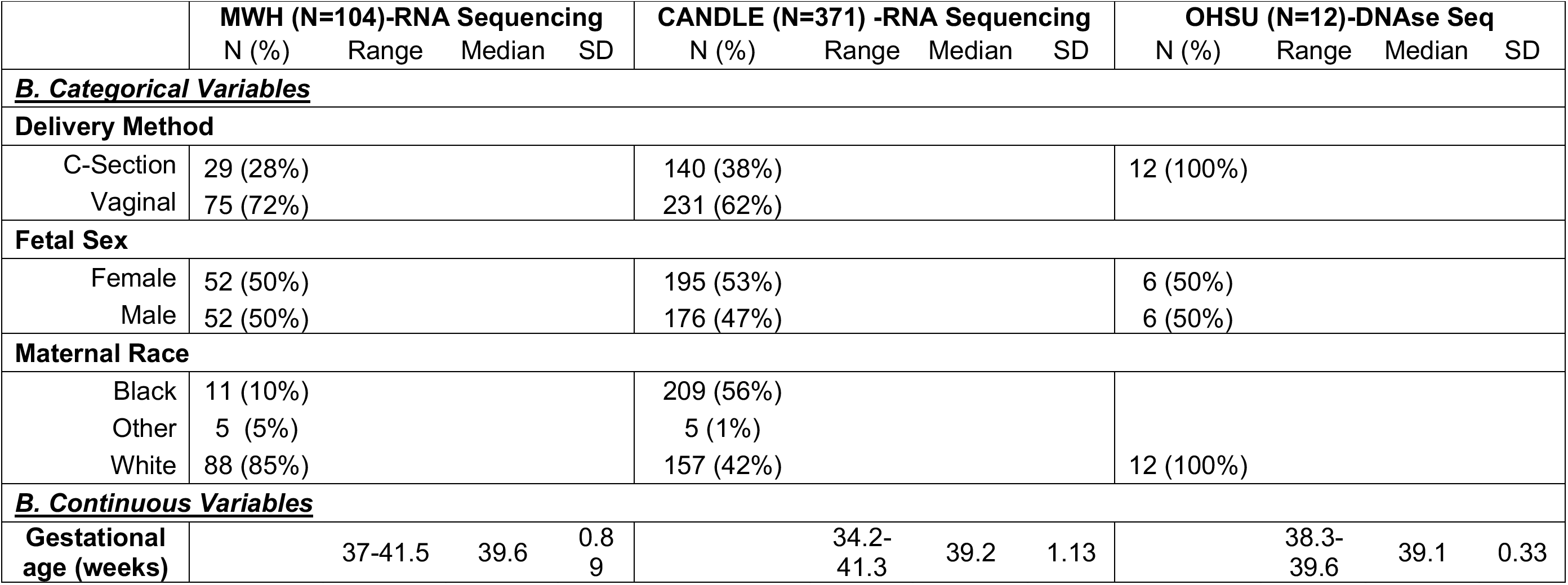
Covariate Data from Cohorts used for RNA sequencing and DNase hypersensitivity Data

|  | **MWH (N=104)-RNA Sequencing** |  |  |  | **CANDLE (N=371) -RNA Sequencing** |  |  |  | **OHSU (N=12)-DNase Seq** |  |  |  |
| --- | --- | --- | --- | --- | --- | --- | --- | --- | --- | --- | --- | --- |
|  | N (%) | Range | Median | SD | N (%) | Range | Median | SD | N (%) | Range | Median | SD |
| **<u>B. Categorical Variables</u>** |  |  |  |  |  |  |  |  |  |  |  |  |
| **Delivery Method** |  |  |  |  |  |  |  |  |  |  |  |  |
| C-Section | 29 (28%) |  |  |  | 140 (38%) |  |  |  | 12 (100%) |  |  |  |
| Vaginal | 75 (72%) |  |  |  | 231 (62%) |  |  |  |  |  |  |  |
| **Fetal Sex** |  |  |  |  |  |  |  |  |  |  |  |  |
| Female | 52 (50%) |  |  |  | 195 (53%) |  |  |  | 6 (50%) |  |  |  |
| Male | 52 (50%) |  |  |  | 176 (47%) |  |  |  | 6 (50%) |  |  |  |
| **Maternal Race** |  |  |  |  |  |  |  |  |  |  |  |  |
| Black | 11 (10%) |  |  |  | 209 (56%) |  |  |  |  |  |  |  |
| Other | 5 (5%) |  |  |  | 5 (1%) |  |  |  |  |  |  |  |
| White | 88 (85%) |  |  |  | 157 (42%) |  |  |  | 12 (100%) |  |  |  |
| **<u>B. Continuous Variables</u>** |  |  |  |  |  |  |  |  |  |  |  |  |
| **Gestational age (weeks)** |  | 37-41.5 | 39.6 | 0.89 |  | 34.2-41.3 | 39.2 | 1.13 |  | 38.3-39.6 | 39.1 | 0.33 |

The model construction process is summarized in **Figure 1,** where we explored the impact of multiple selection in this process. Model construction was done using genomic footprints derived from placental DNAse hypersensitivity data (**Figure 1A**) paired with known TF binding motifs (**Figure 1B**) to generate putative edges between TFs and target genes (**Figure 1C**). Next, we generated a co-expression network of TFs and target genes in our training dataset to parameterize the model (**Figure 1D**). We evaluated different parameters of the model based on how they influenced model size and ability to predict gene expression in our test dataset (**Figure 1E**). We constructed 6 models with different parameters for our training dataset, which are summarized in **Supplemental Table 1**. For the “reference model” (Model 1), we elected to use the median correlation coefficient; then included only the top 15 TFs that were correlated with expression of each target gene in our model. We chose this module size on the basis of analyses performed in the construction of a previous transcriptional regulatory network.^28^ The accuracy of each model was calculated as the R^2^ value of the correlation coefficient (R) between the actual and predicted gene expression in the dataset generated from LASSO regression (out of sample R^2^, or OOSR^2^). The full range of predictions for each individual gene is depicted in **Supplemental Figure S3**. We also assessed the accuracy of the model in our test dataset using random forest classification, but observed minimal differences between the two approaches. For Model 1 (the “reference” model), the out of sample R2 accuracy on the test dataset for each gene from LASSO regression and Random forest were strongly correlated (Cor=0.98, P <2.2×10^-16^, Pearson correlation, see **Supplemental Figure S4**). The OOS R^2^ value for LASSO regression was slightly but significantly higher than random forest in our test dataset (Mean Random Forest OOS R^2^ =0.35, Mean Lasso R2=0.37,P=3.49×10^-7^, t test, **Supplemental Figure S4**). Thus, we concluded that for these samples, LASSO regression slightly outperformed random forest at predicting gene expression in our test dataset.

**Figure 1:**
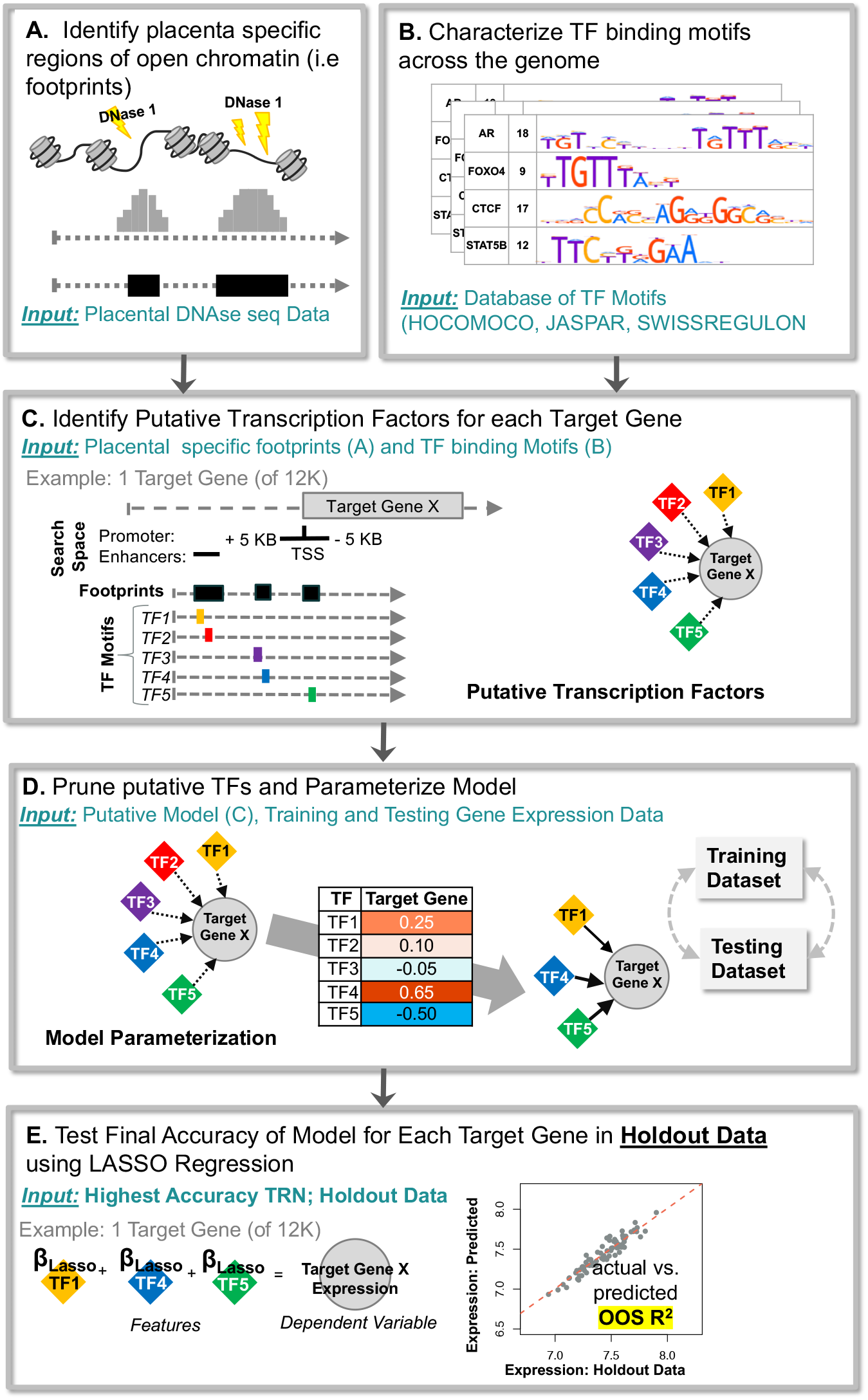
Overview of data and strategy used to construct TRN. First, regions of open chromatin i.e. footprints are identified from Placental DNAse hypersensitivity data (A). Concurrently, TF binding motifs were curated from publicly available databases (B). This data was overlayed (C) to identify putative regulators (TFs) of each target gene. This putative model is pruned using correlations between TFs and target genes established in the training data, and 6 models with different parameters were used. These models were tested in our test data to evaluate the best model (D). The final model was tested in our holdout data (E) to generate final accuracy for each gene and for the overall model

In parallel to our models constructed for TFs with strong correlations in genes with TF binding sites, we also constructed three “null” models; which did not have TF binding sites and/or had weak correlations, and these models are described in **Supplemental Table S2**. We evaluated the accuracy of these models to predict expression in our test dataset using the same approach (**Supplemental Figure S3**). These null models had significantly lower accuracy on predicting gene expression in the test dataset than any of our actual models (P<0.05, Tukey Test, **Supplemental Table S2**). The null models with and without TF binding sites that had correlations between TFs and target genes less than the median training dataset performed the poorest at predicting gene expression in a test dataset. The null models which contained TFs that did not include binding sites at the specific region but were strongly correlated with gene expression performed better than those with poor correlations, but were still significantly less accurate at predicting gene expression in the test dataset than those that also included correct binding data (p<0.05, Tukey Test). Model 9, which contained TFs with no binding sites and poor correlations, had the worst performance for predicting gene expression based on OOS R^2^ in our testing dataset. All analyses did not include the subset of established TFs without motifs as defined by Lambert et al.^29^ We surmised that there were complex regulatory mechanisms by which these TFs act to regulate gene expression (including acting as co-factors). Comparison against the “null” models presented here revealed the extent to which our TRN can predict gene expression.

Overall, the two parameters that altered the model the most were (1) varying the cutoff for the correlation coefficient and (2) the number of TFs included in each model (**Supplemental Figure S3**). Models with a more stringent cutoff contained fewer TFs, target genes and interactions (Models 4 and 5). Model 4 used a higher correlation coefficient cutoff (correlation coefficient >|0.5|), and overall had the highest accuracy of all the models tested (98% of samples had R^2^ >0.25 in test dataset). There was no significant difference in the out of sample R^2^ values between models constructed with and without enhancers (P=1, Tukey Test, see **Supplemental Table 1**). We constructed our model with enhancers to provide complete coverage of potential transcriptional regulators. We elected to construct our model using the parameters established in “Model 3,” using a correlation cutoff of 0.25, as this model was among the more comprehensive in terms of interactions, TFs, and had the highest accuracy outside of Model 4.

### Computational Validation of Model Accuracy

After model parameterization in the training dataset, we then combined the training and testing dataset (N=382) and used this as input data to calculate our final accuracy in the holdout dataset (N=93). This step is critical as a guard against overfitting because it means that the final form of the model was tested once against a holdout dataset that was completely separate from all the variations in the model during the projects development phase,and was selected randomly from samples in the beginning. We analyzed the correlation coefficients in the testing set as well as the ability of the TFs to predict gene expression in the holdout set using LASSO regression (**Figure 2**). In the holdout data, 88% of TF-target gene correlations were statistically significant after adjusting for multiple comparisons (FDR adjusted <0.05), and 83% of TF-target gene relationships were highly correlated with a correlation coefficient of >|0.25 |. Across all genes, LASSO regression using the TFs identified in our model predicted gene expression with a mean OOS R^2^ of 0.39, and we were able to predict 73% of target genes with an OOS R^2^ >0.25. We constructed our TRN using only target genes which we were able to predict with an OOS R^2^>0.25 in our holdout dataset, removing 2,802 genes from our final model.

Our final robust model contained 7,765 genes, with a subset of this model including the 3,213 genes (41.7%) with OOS R^2^ prediction accuracy in our holdout dataset >0.5. The robust model was composed of 113,356 interactions between 391 TFs and 7765 target genes, including 74,014 positively correlated interactions and 38,144 negatively correlated interactions. The median regulon size (target genes per TF) was 156, and was heavily tailed with 75 “hub” TFs which regulated >500 genes (**Figure 3**). Network centrality measures are provided in **Supplemental Table S4.** The 8 most central TFs in our model based on eigenvector centrality (measure of influence of node within a network for each transcription factor) were *SP1, SMAD4, SP3, CTCF, STAT5B, NR2C2, YBX1 and CLOCK*. Our model contained 25 transcription factors that were placenta specific or placenta enriched, based on the Human Protein Atlas^30^, including known placental transcriptional regulators such as *GCM1, DLX3, DLX4*, and *DLX5*. The largest placental enriched transcriptional regulator was *TFAP2A*, which regulated 1,249 different target genes.

**Figure 2:**
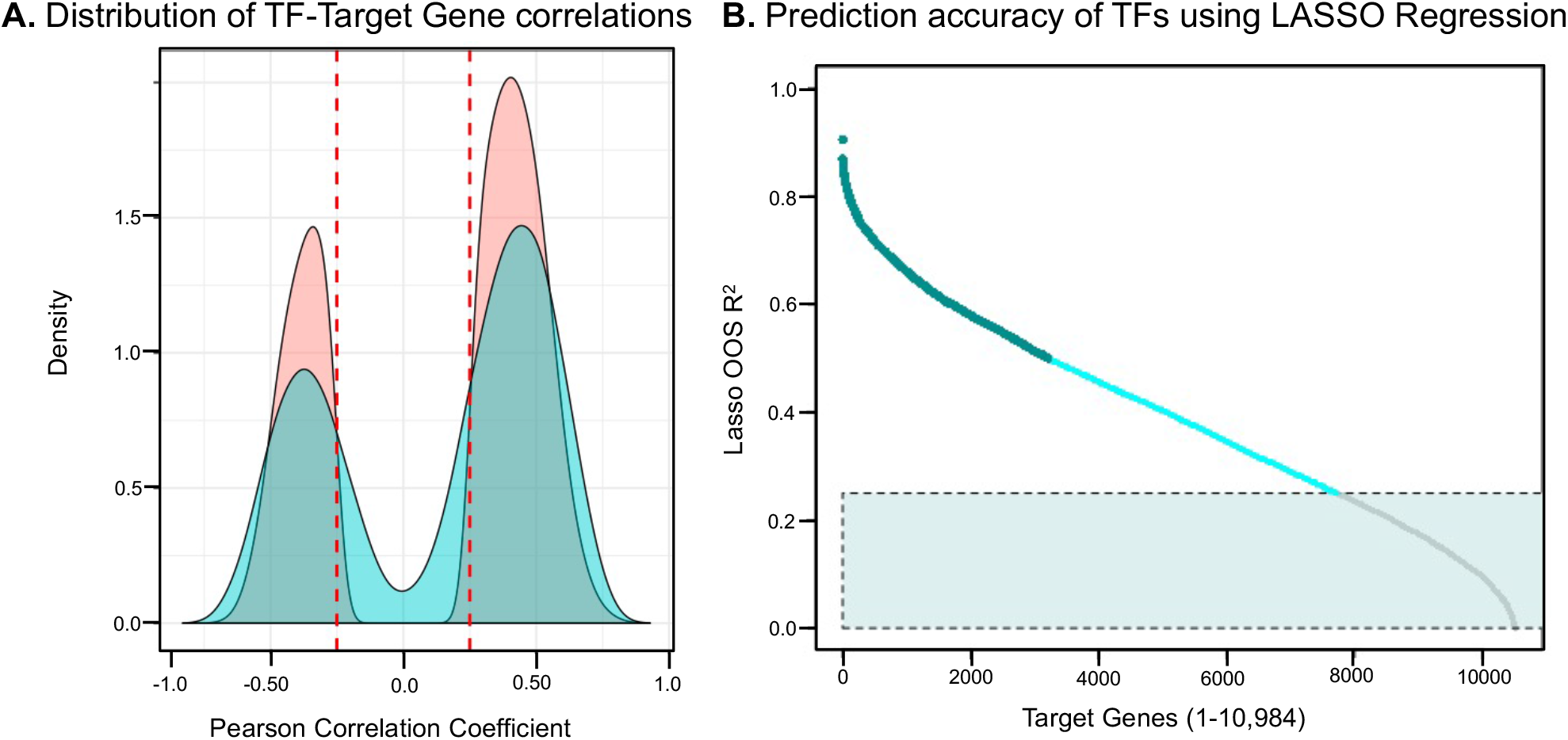
(A.) Distribution of pearson correlation coefficients in training + testing dataset (N=382 samples, pink), and holdout dataset(N=93, blue). Red lines indicate 0.25, which was the cutoff in the training dataset. (B.) Distribution of out of sample accuracy (OOS R^2^) for each of the 10,984 genes calculated in our holdout dataset. Light blue box indicates genes with OOS R^2^ <0.25, which were genes not included in our final model. Dark blue genes were high confidence OOS R^2^.

**Figure 3:**
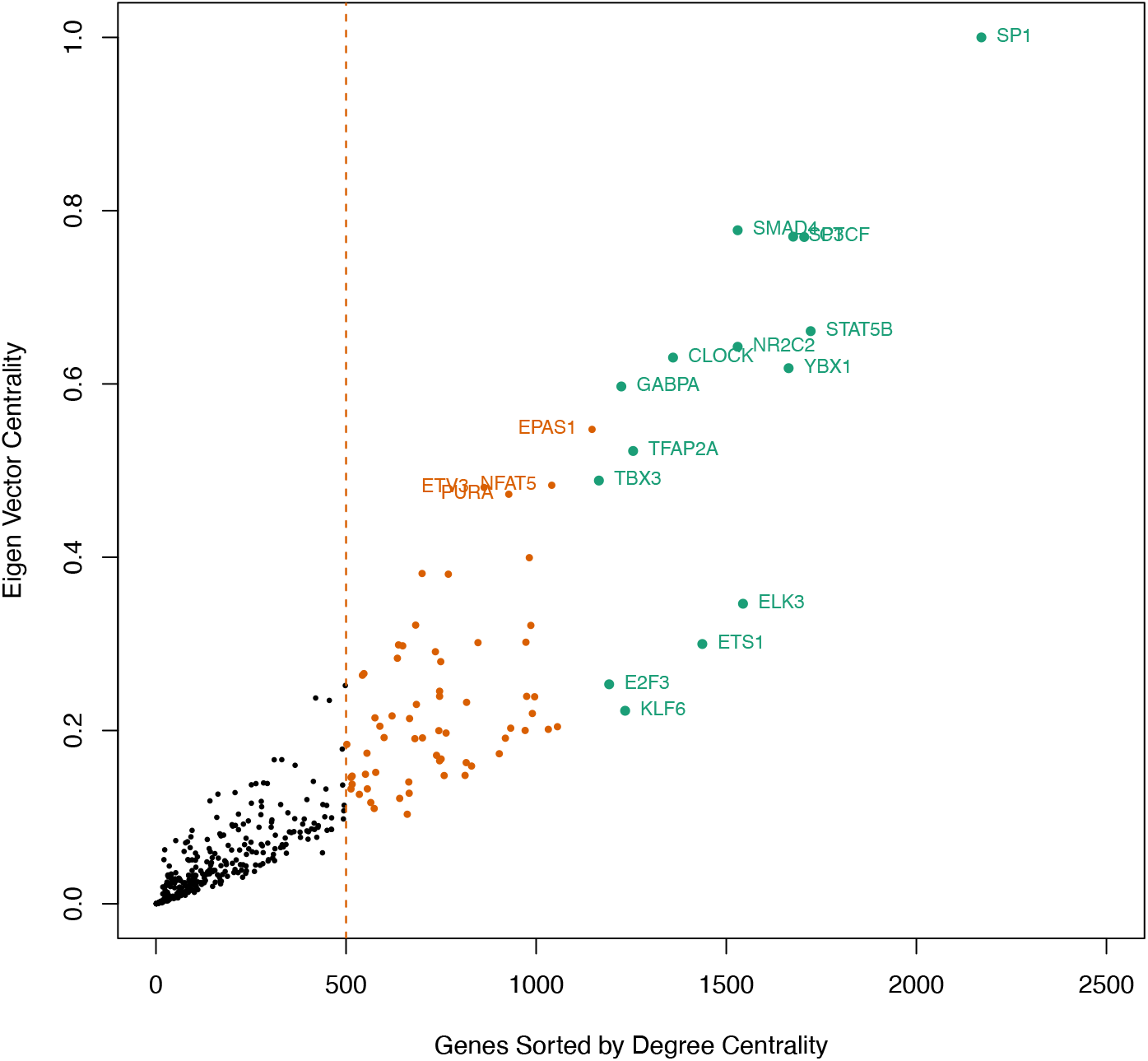
(A.) Summary of network connectivity of 393 TFs based on their target genes. Orange line indicates hub TFs with >500 genes. Top 15 TFs based on degree centrality are shown in green.

### Experimental Validation of Model Accuracy

We performed an siRNA knockdown of four TFs (*GRHL2,CREB3L2, ESRRG, GCM1*) that were selected on the basis of their characterization as “placenta enriched genes” from the Human Protein Atlas ^30^ and which regulated >125 target genes in our model. TFs knockdown was performed in 7 female and 6 male samples (with the exception of *GCM1*, which had 4 male samples, and *GRHL2*, which had 5 male samples). Gene expression for predicted target genes was quantified through RNA sequencing, and we compared the directionality of the knockdown (log fold change of the knockdown vs. control) with the correlation between TF and target gene (**Figure 4**), with the prediction that knockdown of TFs which were positive regulators in the model would result in decreased expression (negative log fold change), and knockdowns of TFs that were negative regulators in the model would result in increased expression (positive log fold change). We achieved an accuracy of 0.96 for *GRHL2*, 0.80 for *GCM1*, 0.71 for ESRRG, and 0.32 for *CREB3L2*. We also analyzed the predictive accuracy and significance of our model within the experimental data in a sex-stratified analysis. In the female siRNA knockdown samples, we achieved a stronger validation of *GRHL2* (Accuracy=1, P=0.015) and *ESRGG* (Accuracy =0.88, P=0.018), and we generally had lower accuracy in male samples (**Supplemental Figure S5**). Overall, our experimental knockdowns revealed that the TRN provided accurate insight into TF regulation for three of our four selected TFs.

**Figure 4:**
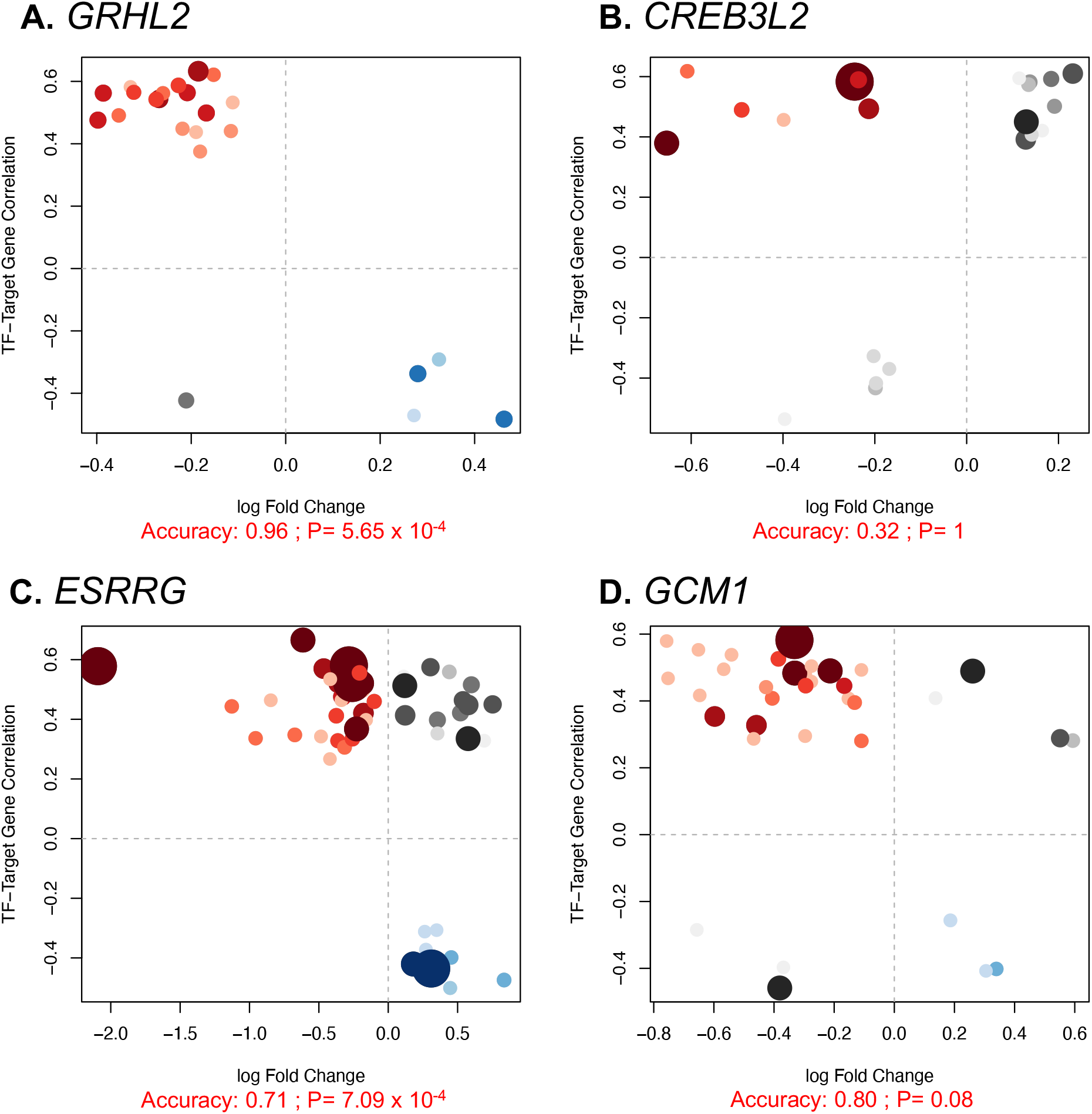
Experimental validation for assessment of final model accuracy. Results of Experimental validation for (A) *GRHL2*, (B) *CREB3L2*, (C) *ESRRG* and (D) *GCM1*. X axis represents the log fold change of knockout vs control, and the Y axis represents correlation between TF and target gene. Model accuracy was calculated as the number of positively regulated genes with a negative log fold change (i.e “true positives”, shown in red) added to the number of negatively regulated target genes with a positive log fold change (i.e “true negatives”, shown in blue) divided by the total number of target genes for each sample. Genes that were not concordant are shaded in grey. The size and shading of each target gene represents the relative rank (1-15) of the TF as a regulator in our TRN.

### Identification of sex-specific differences in transcriptional regulation

Sex specific differences in TF-target gene relationships were captured by generating sex stratified TF-target gene linear models from the complete placental RNA sequencing data (N=475), and comparing the interaction between these 2 linear models. We observed 5,361 TF-target gene relationships between 254 TFs and 1,996 target genes (See **Supplemental Table S5**) that were significantly different between male and females after adjusting for multiple comparisons (FDR adjusted <0.05). Six of these transcription factors and 81 of the target genes were specifically located on the X chromosome, and the rest of the interactions were within autosomal regions. The TFs with the biggest sex specific differences were *ELK3, SP11, ETV5, ETS2*, and *TWIST1*. We compared the difference in estimates between males and females and the direction of the estimate between the TF-target gene relationships and observed four distinct types of sex-specific interactions, as shown in **Figure 5**. 291 positive TF-target gene relationships were stronger in males than in females (Quadrant 1). 3,681 positive TF-target relationships were stronger in females than in males (Quadrant 2). 1261 negative TF-target gene relationships were stronger in females than in males (Quadrant 3), and 120 negative TF-target gene relationships that were stronger in males than in females. (Quadrant 3) The top 3 biggest differences for each quadrant are depicted in **Supplemental Figure S6.** Overall, these perturbed TF-target gene relationships indicate differences in regulation of gene expression between female and male placentas that can be identified in our robust model..

**Figure 5:**
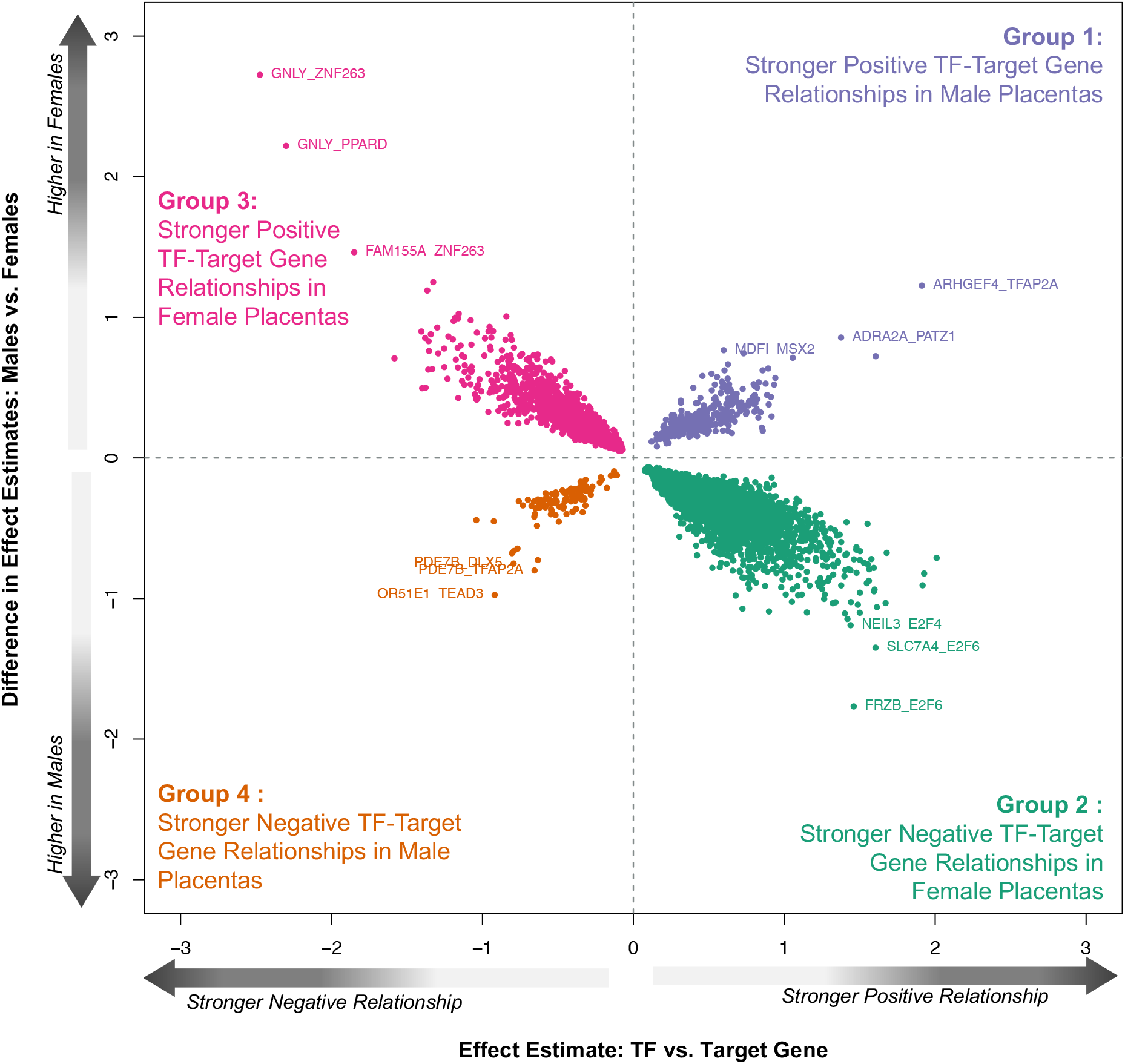
Plot depicting the 5361 sex specific interactions in the placental TRN, divided into four distinct interaction categories. The x-axis represents TF-target gene estimate values from linear regression analysis in the whole dataset. The y-axis represents the difference in estimates between male and female samples.

### Applying the Placental TRN to discover transcriptional regulators of genes associated with preterm birth

Ultimately, the goal of the placental transcriptional regulatory network is to gain insight into transcriptional regulation in the context of prenatal exposures or placental pathologies. As a proof of concept, we used the placental TRN to identify key transcription factors that regulated the group of 129 genes whose placental expression was associated with preterm birth, which we established in an independent publication.^10^ These 129 genes were regulated by a total of 223 transcription factors. We identified enriched transcription factors using hypergeometric tests. Here, the regulons were used in lieu of a gene ontology (GO) or KEGG gene set. Seven TFs (*AHR, ARID5B, BCL6B, NFE2L2, NR3C2, TBX5, THAP5*) were significantly enriched for these pre-term birth associated genes (p<0.1, Fishers exact test). Together, these 7 TFs were responsible for regulation of 35.7% of the differences in expression observed (46/129), including some of our most significantly upregulated (*BIN2*) and downregulated genes (*ADAMTS3*).Analysis of connectivity amongst these TFs showed two clusters (**Figure 6**). *BCL6B* and *TBX5* formed a unique cluster regulating 4 genes together and 2 independent genes each; which were all downregulated in preterm placenta. *BCL6B* was the most statistically significant enriched gene in our network. *TBX5* is classified as a placental-specific gene by the Human Protein Atlas, and it (along with BCL6B) directly regulated 2 other placenta-specific genes (KL and *FAM162B*). The other five enriched TFs (*AHR*, *ARID5B, NFE2L2, NR3C2,THAP5*) formed a highly complex subnetwork that regulated the remaining 38 target genes. Four of these five TFs had identified binding sites and were positively correlated with each other (*AHR, ARID5B, NFE2L2, THAP5*), with *THAP5, AHR, and ARID5B* emerging as the upstream regulators based on hierarchical mapping of the network. Among these genes, *THAP5* and *AHR* had the highest number of differentially expressed genes in total, regulating 20 and 23 genes respectively. This subnetwork revealed that the differentially expressed genes could be attributed to a small number of transcriptional regulators that formed distinct subnetworks.

**Figure 6:**
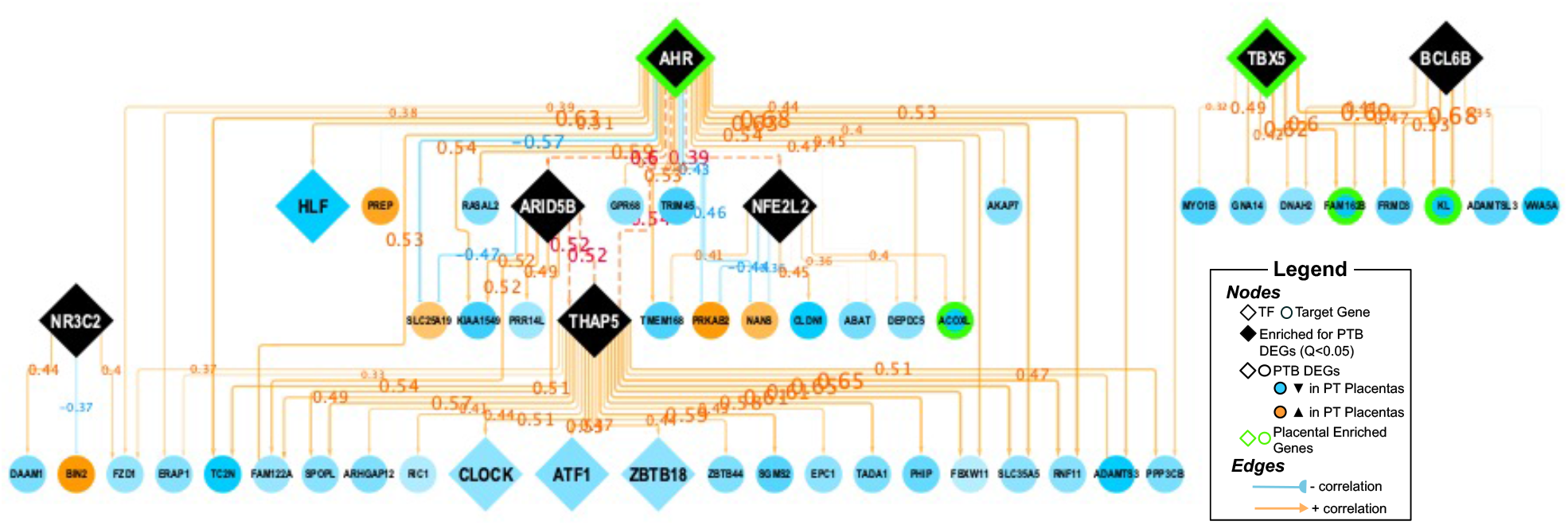
TRN Subnetwork depicting enriched TFs (P<0.1; Fishers Exact Test) (Black) and their corresponding genes that were previously associated with PTB. Color of gene indicates directionality of log fold change in association with preterm birth. Placental specific TFs, defined by the Human Protein Atlas^30^ are outlined in green. Color of edge indicates correlation between TF and target gene, and number represents correlation coefficient. Transcriptional regulation between enriched TFs is highlighted by red dotted edges.

Additionally, six of the 129 differentially expressed genes we previously identified from our previous study^10^ were transcription factors, and these TFs were regulated in part by the enriched TFs. These six TFs themselves regulated downstream genes that were significantly associated with preterm birth (**Table 2)**. *THAP5* was a regulator for two TFs (*ATF1* and *CLOCK1*) that were hub genes in our network and were downregulated in preterm placentas. These transcription factors are crucial for placental transcriptional regulation, and regulate a number of genes in the placenta, including a subset associated with preterm birth. Our TRN revealed that these genes themselves are regulated by a subset of transcription factors enriched for PTB genes. It also highlights that some of the genes previously associated with preterm birth are important to placental transcriptional regulation, based on their degree centrality. These data demonstrate that our placental TRN can be used to identify regulatory pathways driving differential gene expression in placentas from preterm births

**Table 2:** TFs whose placental expression was previously associated with Preterm Birth

| TF | TRN Results |  |  |  | Microarray Results<sup>2</sup> |  |
| --- | --- | --- | --- | --- | --- | --- |
|  | Number of Target Genes | Number of PTB Genes | Genes Previously associated with preterm Birth from microarray<sup>2</sup> | P<sup>1</sup>(Fishers Exact Test) | Log Fold Change | FDR Adusted *P* |
| *ATF1* | 734 | 9 | *DNAH2, FAM122A, TRIM45, EPC1, ZBTB44, TADA1, TC2N, SGMS2, RNF11* | 0.88 | -0.160 | 0.011 |
| *CLOCK* | 1345 | 11 | *RNASEH2C, DAAM1, FIGN, ZBTB44, PPP3CB, RIC1, FBXW11, RASAL2, ARID2, SPOPL, PHIP* | 1.00 | -0.143 | 0.042 |
| *HLF* | 175 | 5 | *ERAP1, SLC25A19, ACOXL, TMEM136, SLC35A5* | 0.17 | -0.270 | 0.011 |
| *IRF2* | 5 | 0 |  | 1.00 | -0.119 | 0.017 |
| *MLX* | 8 | 0 |  | 1.00 | 0.139 | 0.004 |
| *ZBTB18* | 148 | 1 | *ADAMTS3* | 0.92 | -0.144 | 0.035 |
|  | 1. From Fishers Exact Test. 3Log fold change: Preterm vs. Term Samples; FDR adjusted Q, From Paquette, Brockway et al, 2018 |  |  |  |  |  |

## DISCUSSION

In this study, we generated, for the first time, a placenta-specific TRN using established genomic footprinting techniques^31^ and high dimensional RNA sequencing data from 475 placental transcriptomes. Our network captures 113,356 interactions between 391 TFs and 7,765 target genes. Our model was validated against a separate transcriptomic dataset and follow-up experiments for several key regulators. Our study revealed that the TFs *SP1*, *ZNF148,SP3, SMAD4*, and *CLOCK* were master transcriptional regulators of placental gene expression within the TRN; 25 transcription factors unique to the placenta, with TFAP2A emerging as the largest regulon, and a subset of 5,361 TF-target gene interactions that displayed sexual dimorphisms between male and female samples, with more significant transcriptional regulation occurring within female placentas. We applied our model to identify transcriptional regulators of a subset of genes previously associated with preterm birth and found 7 TFs that were enriched as regulators of genes whose placental expression was previously associated with preterm birth, which may indicate clinically modifiable targets. Importantly, this model captures genomic regulation from placentas collected at the end of pregnancy, which is generally when placental samples are readily available and typically collected for large cohort studies. To facilitate additional studies in the community, we have made our model publicly available as a web accessible resource.

### Comparison of the Placental TRN to other Network Models

The major advancement of this study is the generation of a large and sophisticated TRN model using DNAse hypersensitivity data within key placental tissues at term, and the validation of this model using gene expression data captured within a large and diverse cohort. Our model also complements other models of placental transcription. An independent WGCNA of the human placenta constructed using only transcriptomic data revealed 5 hub modules associated with fetal growth restriction (FGR).^24^ One of the core genes of the module associated with both FGR and fetal overgrowth was *CREB3*, which was a hub TF that regulated 738 target genes within our model. Our model builds upon this by inclusion of information about chromatin accessibility using DNAse hypersensitivity data. DNAse hypersensitivity data (without co-expression analysis) has been used to construct a gene regulatory model of early placental transcriptional regulation, including mapping of genes involved in differentiation of human embryonic stem cells to trophoblast cells in mice, and identified 20 key TFs involved in placental differentiation through centrality analysis.^25^ Many of the genes reported as candidates for regulating placental development genes are reported as hub TFs in our model, including *SMAD4, ETS1, RUNX1, CREB1, SMAD3, PPARA*, and *JUND*. Our model captures regulation of key transcription factors identified in other studies from human placental data collected at term. A WGCNA-based analysis revealed that a large proportion of the placental transcriptome is organized into modules of co-expressed genes that were involved in core placental functions and were reproducible across trimesters.^23^ Thus, our model of the placental transcriptome generated at the end of gestation is concordant with previously generated network-based analyses earlier in gestation, suggesting that these TFS are essential not only to placental development but also placental function across gestation.

### Core Transcriptional Regulators of Placental Gene Expression

Many of the highest-ranking core TFs in our model -- including *SP1*, *SP3*, *SMAD4*, and *TFAP2A* – are essential regulators of placental function. *SP1* and *SP3* had the highest eigenvector and degree centrality in our model. SP transcription factors are involved in placental glucocorticoid signaling,^32^ and placental expression of this gene has also been linked to preeclampsia.^33^ *SMAD4* had the second highest eigenvector centrality value, and is a signaling protein in the TGF family which is activated in response to TGF-beta. This signaling pathway plays a crucial role in regulation of trophoblast proliferation, differentiation and invasion, as was reviewed by Adu-Gyamfi et al.^34^ *TFAP2A* regulated 1255 target genes and was the largest placental specific TF based on both eigenvector centrality and degree centrality. *TFAP2A* encodes the activating enhancer binding protein 2-alpha (AP-2α); a transcription factor that is involved in several key processes unique to the placenta.^35^ *TFAP2A* expression is detectable throughout the placenta, including extravillous cytotrophoblasts, cytotrophoblasts and syncytiotrophoblasts.^4^ AP-2α promotes the EGF-dependent invasion of the human trophoblast,^36^ and induces human CRH gene expression.^37^ In our TRN, we were able to recapitulate some of these published findings of *TFAP2A* regulation. In another study, the TFs SP1 and SP3 regulate expression of *TFAP2A* in trophoblast cells.^38^ In our model, *SP1* and SP3 were included in the top 15 TFs regulating the expression of *TFAP2A*. TFAP2A is a positive regulator of CRH in our model; in alignment with work showing that AP-2 induced CRH gene expression.^37^ Thus, our TRN reveals the significance of this TF for placental function as well as some core TFs that regulate many genes within the placenta.

### Sex Specific Regulation of Placental Gene Expression

Sex differences at the molecular level between male and female placentas may contribute in part to differences in growth rates of male and female fetuses and differential responses to perturbations in the in utero environment, and manifestation of disease later in life (fetal programming).^39^ In our analysis, we identified 5,361 TF-target gene relationships that were different between male and female placental samples. Many of these TF-target gene differences may be regulated in part by hormonal differences, as we saw a number of differences related to nuclear hormone receptors that are responsive to sex hormones, including *ESR1, ESSRA, ESRRG*, and *PGR*. We observed a general trend in our interaction terms where both negative and positive TF-target gene relationships were stronger in female placentas compared to male placentas. An aggregated meta-analysis of microarray data revealed 140 differentially expressed genes in male and females samples, with the majority residing in autosomal regions, and identified several TFs that were enriched for these genes, including *NFATC2, AHR, NKX3, HIF1A*, and *REL*.^26^ A number of these TFs were associated in our analysis with sex differences in expression of target genes, such as the *AHR;* which was associated with 111 TF-target gene relationships that were significantly different between males and females. Based on these findings, we hypothesize that there may be stronger transcriptional control of gene expression in female compared to male placentas. These differences may be driven by sex-specific differential response to stress hormones, environmental chemicals, and nutrition, as reviewed by Rosenfeld.^40^ This is significant because there is variation in pregnancy outcomes relate to fetal sex,^41^ such as an increased rate of preterm birth in male infants.^42^ To our knowledge, this analysis is the first to specifically investigate TF-target gene differences by sex within the context of a TRN. Further analysis needs to be performed on these relationships, including factors that modify these relationships and the role of genomic imprinting.

### The Placental TRN reveals Transcriptional Regulators of Prematurity

Our ultimate goal is to apply this placental TRN as a tool to help identify perturbed regulation in advance pregnancy outcomes, or in relation to environmental exposures. To this end, we used the TRN to identify seven different transcription factors enriched for genes associated with preterm birth that we identified in an independent study.^10^ *TBX5* and *BCL6B* regulated a distinctive group of PTB genes within our TRN; and *BCL6B* was the most enriched TF in our model (p=0.016). *BCBL6B* encodes the B-cell CLL/lymphoma 6 member B protein, a transcriptional repressor which mediates VEGF signaling and angiogenesis,^43^ which are both key to placental function. *TBX5* has been previously linked to PTB,^44^ and interestingly is placentally enriched in the human protein atlas,^30^ suggesting potential function in placental development and maturation. The other 5 TFs were highly co-regulated within our model, with *THAP5, ARID5B* and *AHR* emerging as the master regulators within this subnetwork. *THAP5* encodes a transcriptional repressor that is not well studied due to a lack of orthologs in mice and humans, but may have pro-apoptotic functions.^45^ *ARID5* encodes AT-rich interactive domain-containing protein 5B which is a transcriptional coactivator involved in cellular growth and differentiation. Methylation of the *ARID5B* promoter region in the placenta has also been associated with decreased fetal birthweight.^46^

Two of the TFs that were enriched for preterm birth associated genes, *AHR* and *NFE2L2*, are important xenobiotic response genes that respond to environmental perturbations and are linked to inflammation AHR encodes the Aryl Hydrocarbon Receptor, a nuclear hormone receptor which is activated by exogeneous ligands. Beyond its role in cellular detoxification, this receptor is also involved in cellular proliferation, migration and development,^47^ and regulation of the innate and adaptive immune system.^48^ *AHR* is highly expressed throughout pregnancy, and protein expression increases five-fold between first trimester and term placenta within syncytiotrophoblasts of chorionic villi.^49^ AHR agonists alter the vascular remodeling of the placenta, resulting in reduced volume of the placental labyrinth zone.^50^ AHR is hypothesized to be involved in preterm birth through promotion of pro-inflammatory cytokines.^51^ Prenatal exposure to AHR agonist TCDD increases the incidence of preterm birth in mice^52^ and induces pro-inflammatory responses in human placental explants.^53^ Placental inhibition of AHR has been shown to inhibit cell migration as a potential mechanism.^54^ Specific AHR agonists such as cigarette smoke are associated with increased odds of developing preterm birth.^55^ In our model, *AHR* regulated expression of Nuclear factor erythroid 2-related factor 2 (NRF-2; encoded by *NFE2L2*), which was also enriched for preterm birth genes. There is substantial cross talk between NF-2 and AHR,^56^ and our work is concordant with previous analyses demonstrating that AHR regulates NRF-2 expression by binding to the promoter region of the gene in other tissues.^57^ We observed this same relationship within our own TRN as AHR was one of the significant positive regulators of NRF-2 within the placenta. NRF-2 is a master regulator of antioxidant response and its products protect cells from oxidative insults.^58^ NRF-2 has been implicated as a modulator of oxidative stress in early pregnancy.^59^ NRF-2 deletion in mice leads to increased inflammation and premature labor after treatment with LPS, changes to the volume of the labyrinth zone of the placenta, reduction in placental volume, increased oxidative stress, elevated inflammatory cytokines and cell death.^60,61^ In human studies, alterations in NRF-2 are attributed to changes in placental cells and preeclampsia.^60^ Our TRN reveals that many of the preterm birth associated genes are regulated by AHR and NRF-2, which could indicate activation of these gene expression pathways in preterm placentas.

### Applications of the Placental TRN

Placental omics data can be used to quantify perturbations that influence gene regulation and physiological activity, providing insight into the underlying etiology related to outcomes or exposures.^62^ Many researchers studying the placenta have used omics data to generate lists of genes which are associated with prenatal exposures as well as infant or child health outcomes, as summarized in our recent review.^63^ These genes are often contextualized using pathway analysis approaches within KEGG or Gene Ontology (GO) gene sets. The TF enrichment analysis we have performed using regulons is complementary to using other defined gene sets such as KEGG or GO. TRNs are focused on the question: *“What transcription factors regulate these genes within the placenta?”* There are other applications that provide an opportunity to query TF binding, sites such as ChEA^64^, which are based on aggregation of Chip-seq data that is not tissue or cell type specific. Our TRN provides a placental tissue-specific version that can be specifically queried to investigate individual TFs or target genes of interest; or to perform an enrichment analysis similar to what we have performed using genes associated with premature birth. We have presented the full results of the genome-scale model in the format of a searchable web tool, PlacentalTRN, which is available at https://github.com/AlisonPaquette/PlacentalTRNRShinyApp. This model has 3 features: (1) a target gene search feature, (2) a transcription factor search feature, and (3) an enrichment test. The search tools allow users to search the model for specific genes or specific transcription factors, and evaluate model accuracy and predicted targets on a case-by-case basis. The enrichment test tool allows users to import a list of genes (such as the list of preterm birth genes presented here), and identify transcription factors which regulate these genes. For those interested in accessing the TRN independently of the web interface, we provide the entire TRN in **Supplemental Table 6**. Our intention is that other placental biology researchers will be able to easily access the TRN and identify transcriptional regulators of their own gene lists of interest by using this database.

### Strengths and Limitations

Our model of placental transcriptional regulation should be interpreted with limitations in mind. We use TF expression as a proxy for TF activity, which does not account for post transcriptional or post translational modifications, and may be confounded by temporal and spatial disconnects, cellular heterogeneity, and differences in proliferation,^65^ which are common for all studies that employ this approach. We anticipate confounding due to cellular heterogeneity, as TRNs reflect epigenetic regulation dependent on regions of open chromatin and DNA methylation which is intrinsically cell but not tissue-type specific, and our results are not built on placental data which is a composite of different cell types. This can create false positive relationships and dilute signal strengths between known TFs and target genes. For each target gene, we present only the top 15 TFs that are best predicted to regulate expression. Thus, we may not represent all the TFs with binding sites within that region. Our analyses are composed of data solely from placentas at the end of gestation from relatively non-pathological pregnancies without preeclampsia, and so may be inappropriate for researchers studying earlier developmental time points, and also may not represent perturbed TF-target gene interactions related to those pathologies. As always, it will be important to further test the model in an entirely separate validation cohort collected from different participants as they become available. In the future, we hope to build upon this model by including cell type specific interactions and temporal resolution of time series changes that occur in the placental transcriptome.

Our model makes a contribution as the first publicly available placental transcriptional regulatory network, as well as one of the first models of transcriptional regulation of any organ with an accessible and searchable database that allows researchers to perform their own analysis. We applied rigorous methods to test the accuracy of our model, including an independent validation dataset which is the gold standard in machine learning approach.^66,67^ To our knowledge, our model is composed of the largest placental transcriptomic dataset constructed to date. We also provide an innovative assessment of sex differences in TF-target gene relationships in our model, which further captures the importance of fetal sex as a biological variable in placental omics research. Finally, we have tested our TRN using isolated villous cytotrophoblasts collected from placentas of non-pathological women delivering at term, which are more reflective of the term placental transcriptome than commercial placental cell lines which are commonly used for transcriptomic analyses. This data could potentially be used in the future to shed insight into transcriptional mechanisms related to pregnancy complications, and identify pharmacological interventions to target specific transcription factors that alter placental function.

## MATERIALS AND METHODS

### 1. Sample Information

To construct the TRN, we integrated transcriptomic and chromatin accessibility data from placental tissue generated from 3 different sources as described below.

#### 1A. Magee Women’s Research Institute (MWRI) Samples

Tissue was collected from non-pathological placentas from pregnant individuals who delivered at the Magee Womens Hospital in Pittsburgh, PA. Inclusion criteria included: Maternal age >18 years, signed informed consent, singleton pregnancies and gestational age >37 weeks.

Exclusion criteria included pregnancy complications including, fetal growth restriction (FGR), chromosomal abnormalities, preeclampsia, gestational diabetes, chorioamnionitis, antepartum hemorrhage, and a previous history of preterm birth. Placental biopsies (5 mm^3^) were sharply dissected within 30-40 minutes after delivery using a region of the placenta that is midway between the cord insertion and the placental margin, and between the chorionic and basal plates, and free of any macroscopic lesions, as we previously detailed.^68^ Tissue biopsies were placed in RNAlater for 48 h in 4°C, then stored in −80°C until processed for RNA sequencing. Directional RNA-seq was performed by the Genomics, Epigenomics and Sequencing Core (GES Core) at the University of Cincinnati. Approximately 50 mg tissue was homogenized in 0.8 ml Lysis/Binding Buffer from mirVana miRNA Isolation Kit (Thermo Fisher, Grand Island, NY) by using FastPrep-24 5G homogenizer (MP Biomedicals, Solon, OH). Total RNA extraction was conducted according to manufacturer’s protocol. RNA quality was examined on an Agilent 2100 Bioanalyzer (Santa Clara CA). rRNA was depleted using the Ribo-Zero Gold kit (Illumina, San Diego, CA) with 1 μg of total RNA as input for the SMARTer Apollo NGS library prep system (Takara, Mountain View, CA). The library for RNA-seq was prepared by using NEBNext Ultra II Directional RNA Library Prep kit (New England BioLabs, Ipswich, MA) and the sequencing was run on an Illumina HighSeq2000 with paired end reads, 101bp read length, and at 50 million read depth.

#### 1B. CANDLE Samples

Placental tissue was collected from 371 non-pathological placental transcriptomes collected from women from Shelby County Tennessee as part of the CANDLE (Conditions Affecting Neurocognitive Development and Learning in Early childhood) study. Inclusion criteria included: Maternal age >18 years with signed informed consent. Exclusion criteria included infants categorized as having severe pregnancy complications recorded in the CANDLE medical records, including confirmed clinical chorioamnionitis, preeclampsia, oligohydramnios, placental abruption, infraction or previa, and fetal chromosomal abnormalities. Methods of placental sample collection and RNA sequencing processing for this cohort have been previously described.^69^ Briefly, within 15 minutes of delivery, a piece of placental villous tissue approximately 2 x 0.5 x 0.5 cm was dissected from the placental parenchyma. The tissue cubes were stored in 20 ml RNAlater and refrigerated at 4°C overnight (at least 8 hours but no more than 24 hours). Following this, each cube was transferred to an individual cryovial containing fresh RNAlater and stored at −80°C. The fetal villous tissue was manually dissected and cleared of maternal decidua. RNA was isolated using the AllPrep DNA/RNA/miRNA Universal Kit (Qiagen; Germantown, MD) according to the manufacturer’s recommended protocol. RNA purity was assessed by measuring OD_260/230_ and OD_260/260_ ratios with a NanoDrop 8000 Spectrophotometer (Thermo Fischer Scientific; Waltham MA). RNA integrity was determined with a Bioanalyzer 2100 using RNA 6000 Nanochips (Agilent; Santa Clara, CA). cDNA libraries were prepared from 1 μg of total RNA using the TruSeq Stranded mRNA kit (Illumina, San Diego, CA) and the Sciclone NGSx Workstation (Perkin Elmer, Waltham, MA). Prior to cDNA library construction, ribosomal RNA was removed by means of poly-A enrichment. Each library was uniquely barcoded and subsequently amplified using a total of 13 cycles of PCR. Library concentrations were quantified using Qubit fluorometric quantitation (Life Technologies, Carlsbad, CA). Average fragment size and overall quality were evaluated with the DNA1000 assay on an Agilent 2100 Bioanalyzer. Each library was sequenced to approximately a depth of 30 million reads on an Illumina HiSeq sequencer.

#### 1C. Oregon Health Sciences Center Placental Repository (OHSU)

12 placental villi samples were derived from non-pathological, pregnancies from pregnant individuals delivering via caesarean section at the OHSU Hospital labor and Delivery Unit. Samples were specifically selected to be equally matched across fetal sexes, and from term placentas (>37 weeks). Exclusion criteria included maternal BMI >25, maternal age <18 or >40, any current pregnancy complications and current smokers. Placentas were collected and weighed immediately following cesarean section. Five random samples of tissue (~80 grams) were collected from each placenta and placed in PBS to be transported back to the lab. The chorionic plate and decidua were removed from each randomly isolated placental sample, leaving only villous tissue, which was thoroughly rinsed in PBS to remove excess blood. Primary cytotrophoblast were isolated from villous tissue using a protocol adapted from Eis *et al*. using trypsin/DNase digestion followed by density gradient purification. Isolated cytotrophoblast cells (5-10×10^6^ cells /ml) were then frozen in freezing media at (10% DMSO in FBS) and stored in liquid nitrogen until usage. Placentas were collected f into a tissue repository under a protocol approved by the Oregon Health & Sciences University Institutional Review Board with informed consent from the patients. All tissues and clinical data were de-identified before being made available to the investigative team.

### 2. Quantifying Placental Gene Expression in the CANDLE AND MWRI Placentas

A flowchart describing sample preprocessing is described in Supplemental **Figure S1B**. For the RNA sequencing samples from MWRI and the CANDLE cohort, transcript abundances were estimated using the quantification program Kallisto.^71^ Initially quality control was performed on each RNA sequencing sample using FastQC.^72^ Length scaled counts were condensed to Ensembl Gene IDs using the Bioconductor TXImport package.^73^ Genes were then filtered to remove low expressing genes using the “FilterByExprs” function of EdgeR, which by default removes genes with <10 counts (0.5 CPM) in 70% of samples.^74^ For the dataset from MWRI, this included 17,574 genes, and from the CANDLE samples, this included 16,440 genes. Datasets were then normalized using the trimmed mean of M,^75^ and then precision weights were generated for log-cpm values using the voom transformation.^76^ The datasets were then merged by all genes common across both datasets, generating a final transcriptomic dataset of 14,837 genes in 475 samples. Confounding related to both RNA sequencing batch and intrinsic differences between the cohorts was eliminated using the comBat algorithm from the Bioconductor SVA package,^77^ which uses an empirical Bayes method to estimate location and scale parameters for each batch, which are then removed to provide a consistent mean and variance for each gene, across all batches. This approach has successfully been used to integrate microarray datasets,^10^ as well as RNA sequencing datasets.^78^ RNA sequencing batch was set as the “batch” variable to be adjusted for (The CANDLE cohort was sequenced in 4 batches, and the MWRI dataset was sequenced as one batch), and maternal race and gestational age were included as covariate variables in the model matrix, to ensure that differences related to these variables would be retained in the final dataset. Batch effects were evaluated using principal components analysis (See **Supplemental Figure S2; Supplemental Table 1).** For TRN Construction, the data was split into a holdout dataset (20% of samples-N=93), and 20% of the remaining samples was used for the testing dataset (N=75), with 80% used for the construction of the initial TRN (N=307). After model splitting but prior to model construction, we tested to ensure that there were no statistical differences between the training and holdout datasets in relation to the distribution of covariates including batch/cohort, labor status maternal race, and fetal sex, (categorical, tested using chi squared tests) as well as gestational age and birthweight (continuous, T-tests). In the final model, all samples were used (N=475 samples).

### 3. DNase Hypersensitivity Data Generation

DNase hypersensitivity data was generated from the samples collected at OHSU. Altius Institute used an established protocol for tissue processing developed as part of the ENCODE project.^79^ Samples were processed using the DHS-pseudonano assay & nuclei were sterile filtered (0.25-6.4 million nuclei/sample), with minimal cellular debris, and treated with 3 doses of DNase (40, 60 and 80 units) followed by a stop buffer including RNAse A, followed by treatment with proteinase K. The samples had a spot (signal portion of tags) score >0.5 except for 1 sample (which had a score of 0.34), which is a measure of enrichment, reflecting the fraction of reads that fall within peak calling regions. A score >0.4 is indicative of high quality DNase sequencing data meeting ENCODE quality guidelines. Peak regions of DNase-seq data were identified using F-seq,^80^ a density estimator for high-throughput sequence tags, using a threshold parameter of 2.5, which was selected based on previously reported performance metrics.^81^ The peak regions and DNase-seq data were both used to detect genomic “footprints”–i.e regions of accessible–DNA using HINT,^82^ a method based on hidden Markov models.

### 4. Transcription Factor Motifs & Single Gene Model Construction

For each gene in our network, we identified putative TFs through integration of the DNAse hypersensitivity data and TF binding motif information. Transcription factors were defined by the HumanTFDB which had motifs from these the TF databases (667 TFs).^29^ Transcription factor motifs were identified using three established databases: JASPAR,^83^ HOCOMOCO,^84^and Swissregulon.^85^ We first matched these binding sites with the footprints in the placenta (described above) using FIMO (Find Individual Motif occurrences), applying the default 10-4 match threshold (Version 5.0.5).^86^ The motifs were aggregated into position weight matrices within the R package “MotifDB”.^87^ We assigned motifs to their TFs as well as those in the same DNA-binding domain family using the TFClass database, as established in our previous work.^17^ In our analysis, we did not consider the subset of transcription factors classified by Lambert et al with no known motifs. We included both promoter and enhancer regions in our model. The promoter region was defined as +/− 5KB from the TSS which has been shown to maximize target gene prediction in transcriptional regulatory networks constructed with footprinting data.^88^ The Enhancer regions were identified using genehancer (Version-4.11).^89^ Genehancer is a database that integrates reported enhancers from multiple sources, including ENCODE and Ensembl, and uses a combinatorial likelihood method to define genes based on tissue co-expression correlations, enhancer-targeted transcription factors, expression quantitative trait loci, and chromatin conformation capture. We included all enhancer regions defined by Genehancer as “elite”, indicating that they had more than 2 evidence sources, and those defined as “placental-specific” based on Genehancer analysis of tissue specific genes.

### 5. TRN Construction & Parameterization

#### 5A. Constructing and evaluating Model in Training Data

We integrated the data described above to generate a model consisting of nodes and edges. A “putative edge” was identified between a TF and a target gene if a motif for that transcription factor was present in a region of open chromatin (i.e footprint) in the gene’s promoter or enhancer (see Figure 1). We generated a co-expression matrix for each “putative edge”. We ranked each target gene based on the absolute value of the pearson correlation coefficient. We pruned our model through an initial “baseline model” (Model 1), including only TFs with > median absolute value of the overall Pearson correlation coefficient for each target gene, then including only the top 15 genes based on absolute Pearson correlation coefficient. We elected this initial module size of 15 based on similar analyses performed in the construction of a transcriptional regulatory network constructed in the context of T-helper Cell Type 17 (Th17) differentiation.^28^ To evaluate the accuracy of our initial TRN and select model parameters for our full model, we used LASSO regression and random forest to evaluate how the expression of the TFs in our model predicted gene expression of each target gene in our testing dataset. The R^2^ value was calculated as the squared estimate of the correlation between the predicted vs. actual expression values. We selected the best model to use in the holdout dataset based on these thresholds. Machine learning was performed using the “glmnet” (version 2.0-16) and “random forest’ (version 4.6-14) R packages.

#### 5B. Null Model

We created three “null” TRNs to evaluate the validity of our model to distinguish between signal and noise, which we constructed in our training and evaluated in testing dataset. The first null model was constructed from 15 randomly selected genes with motifs but with correlations that were less than the median correlation coefficient. In parallel to our model constructed above, we generated a network consisting of “null putative edges” defined as interactions between a TF and a target gene if a motif for that transcription factor was not present in a region of open chromatin (i.e, footprint) in the genes promoter or enhancer. We generated a co-expression matrix for each “null putative edge” within our training dataset. Our second null model was constructed of 15 randomly selected genes with no motifs but with correlation coefficient’s > the median value in our true dataset. Our third null model was constructed of 15 randomly selected genes with correlation coefficients less than the median value in our true dataset and with no motifs. In some instances, 15 genes were not available that met these criteria, and in this case, we included all the genes available.

#### 5C. Final Model Construction

After initial model parameterization in our training and testing dataset, we moved on to combine these 2 datasets, and then test the model in our final holdout dataset using the best parameters as identified from our original models. We did not include target genes with an out of sample R^2^ accuracy <0.25. In the full model, network connectivity scores were calculated using the R package “igraph” (Version 1.2.4.1). To identify sex-specific differences in TF-target gene relationships, we ran linear regression analyses for each target gene-TF interaction in our TRN with fetal sex as an interaction term. Models were considered statistically significantly different by sex if the FDR adjusted Q of the interaction term was <0.05. We ran stratified models in male and female samples to calculate the coefficient estimate and identify differences in these estimates.

### 6. Experimental Validation

We performed knock downs of selected TFs in primary villous trophoblast cells from placental samples collected through the OHSU placental biorepository. Placenta specific TFs were identified using the Human Protein atlas ^30^, then 4 TFs were selected based on module size (>125 target genes) and median gene expression (> Median RPM). Separate transfections were performed using the ON-Target SMARTpool siRNAs (Dharmacon) to induce silencing of four genes: *CREB2L2:* (Catalog # L-019357-02-0005), *ESRRG* (Catalog # L-003405-00-0005), *GCM1* (Catalog # L-011491-00-0005) and *GRHL2* (Catalog # L-014515-02-0005) with a non-targeting negative control (Catalog # D-001810-10-05). siRNAs were diluted in nuclease free water and stored at a 50 μM stock concentration. These transfections were each performed from villous trophoblasts collected from elective c-sections at OHSU as described above.

For transfection, cytotrophoblast cells were rapidly thawed in a 37°C water bath and immediately diluted in Iscove’s modified Dulbecco’s medium (25mM glucose, 4mM glutamine and 1mM pyruvate (Gibco #12440-046) supplemented with 10% FBS (Corning #35-010-CV) and 1% penicillin/streptomycin (Gibco #15140-122) termed as complete media. Cells were centrifuged at 1000g for 10 min and suspended in fresh complete media. The cells were plated at 2×10^6^ cells per well concentration in 12-well culture plates and cultured at 37°C, 5% CO_2_. After 24 hours, cells were transiently transfected using the above siRNAs and Lipofectamine (Invitrogen #11668-019) as per manufacturer’s protocol and incubated for another 24hrs. At the end of 24hrs (total of 48hrs from the start of culture) the transfection media was replaced with complete media and cells were cultured for another 24hrs. After 72h of total culture time (24h following previous complete media change) the cells were lysed in 700uL of Qiazol and RNA was isolated using the Zymo Direct-zol Mini-prep kit and reagents (Zymo-Catalog # RD2051). RNA integrity was determined with a Bioanalyzer and only RNA samples with a RNA Integrity Number (RIN) > 8.6 were used for RNA-Seq analysis. cDNA libraries were prepared from 1 μg of total RNA using the Illumina Stranded Total RNA library kit. Each library was sequenced to approximately a depth of 50 million reads on an Illumina NovaSeq 6000 S4 (2 x100 read length) sequencer by the OHSU sequencing core.

We used a similar pipeline to process the RNA sequencing data as was used for the human placental data, as described above. Transcript abundances were estimated using Kallisto, and length scaled counts were condensed to Ensembl Gene IDs using the Bioconductor TXImport package.^73^ Genes were then filtered to remove low expressing genes using the “FilterByExprs” function of EdgeR.^74^ Differentially expressed genes between siRNA knockdowns vs. negative scrambled controls were identified using generalized linear models implemented within EdgeR,^74^ including only target genes that were predicted to be expressed by each TF. Dispersion parameters were estimated using the Cox-Reid method, and then differentially expressed genes were identified using the using quasi-likelihood F-tests, which is more robust and produces more reliable error rates when the number of replicates is small.^90^ We analyzed only genes that were statistically significant using an unadjusted p<0.05. Model accuracy was calculated at the number of positively regulated genes with a negative log fold change (i.e “true positives”) added to the number of negatively regulated target genes with a positive log fold change (i.e “true negatives”) divided by the total number of target genes for each sample. Statistical significance was assessed using a one sided Fishers exact test on the number of genes that were “true positives” and “true negatives” versus other genes.

### 7. Enrichment of TF Modules associated with Preterm Birth

Differentially expressed genes significantly associated with preterm birth were identified through ancillary analyses of aggregated microarray data.^10^ In this prior analysis, genes were considered statistically significantly associated with preterm birth using an FDR adjusted Q value <0.05. We performed an enrichment analysis^91^ in which we considered the transcription factors and all the genes they regulate (i.e regulons) as gene groups (analogous to predefined pathways such as KEGG or GO), and compared if the list of differentially expressed genes were enriched within specific regulons using one sided Fisher’s exact test. We considered the 7712 target genes captured within our TRN as the background genes for analysis. We only included TFs in our enrichment tests which regulated three or more differentially expressed genes. We considered a TF to be a key regulator if its target genes were over-represented based on a P value <0.05.

All code is publicly available in the following github repository (https://github.com/AlisonPaquette/PlacentalTRN_ManuscriptCode/). All analyses were performed in R (Version 3.6.1), and data was visualized using Cytoscape (Version 3.7.2). The TRN was developed using the R Shiny Web Application.

## ACKNOWLEGMENTS

We would like to acknowledge all the cohorts and biorepositories that provided placental tissue samples, including participants and research staff and investigators who made this possible. The samples from Magee Women’s Research Institute were collected by the Perinatal Biology Lab (PI, YS) from specimens by the Obstetrical Specimens Procurement Unit at Magee Women’s Hospital, based on published protocols, detailed in methods. The authors wish to thank our ECHO colleagues, the medical, nursing and program staff, as well as the children and families participating in the ECHO cohorts. We also acknowledge the contributions of the following ECHO program collaborators: 1. Coordinating Center: Duke Clinical Research Institute, Durham, North Carolina: Smith PB, Newby KL, Benjamin DK. 2. Data Analysis Center: Johns Hopkins University Bloomberg School of Public Health, Baltimore, Maryland: Jacobson LP; Research Triangle Institute, Durham, North Carolina: Parker CB.

## Disclaimer

Current Address for Dr. Brockway is the Center for Scientific Review, National Institutes of Health, Bethesda Maryland, United States. This article was prepared while Dr. Heather Brockway was employed by the University of Florida. The opinions expressed in this article are the author’s own and do not reflect the view of the National Institutes of Health, the Department of Health and Human Services, or the United States government.

## Author contributions

Conceptualization: AGP, LM, LM, SS, NP
Methodology: YMW, PS, JP, CF, JM, TB, PB
Software: PS, CF, YMW, RR
Investigation: KA, YMW, JP, HL, MB, LK, SH, HB
Resources: SS, CK, WAM, NB, KL, YS, JS
Visualization: AGP, RR
Supervision: NP, SS, LM, YS, HJ, LM
Writing—original draft: AGP, NP
Writing—review & editing: AGP, KA,YMW, JP, HL, PS, MB, LK, SL, RR, CF, JM, TB, PB, HB, WAM, NB, KJ, CK, JS, LM, HJ, YS, LM, SS, NP

