## Supplemental Data 7 for "A Genome Scale Transcriptional Regulatory Model of the Human Placenta"

**SUPPLEMENTAL FIGURES**

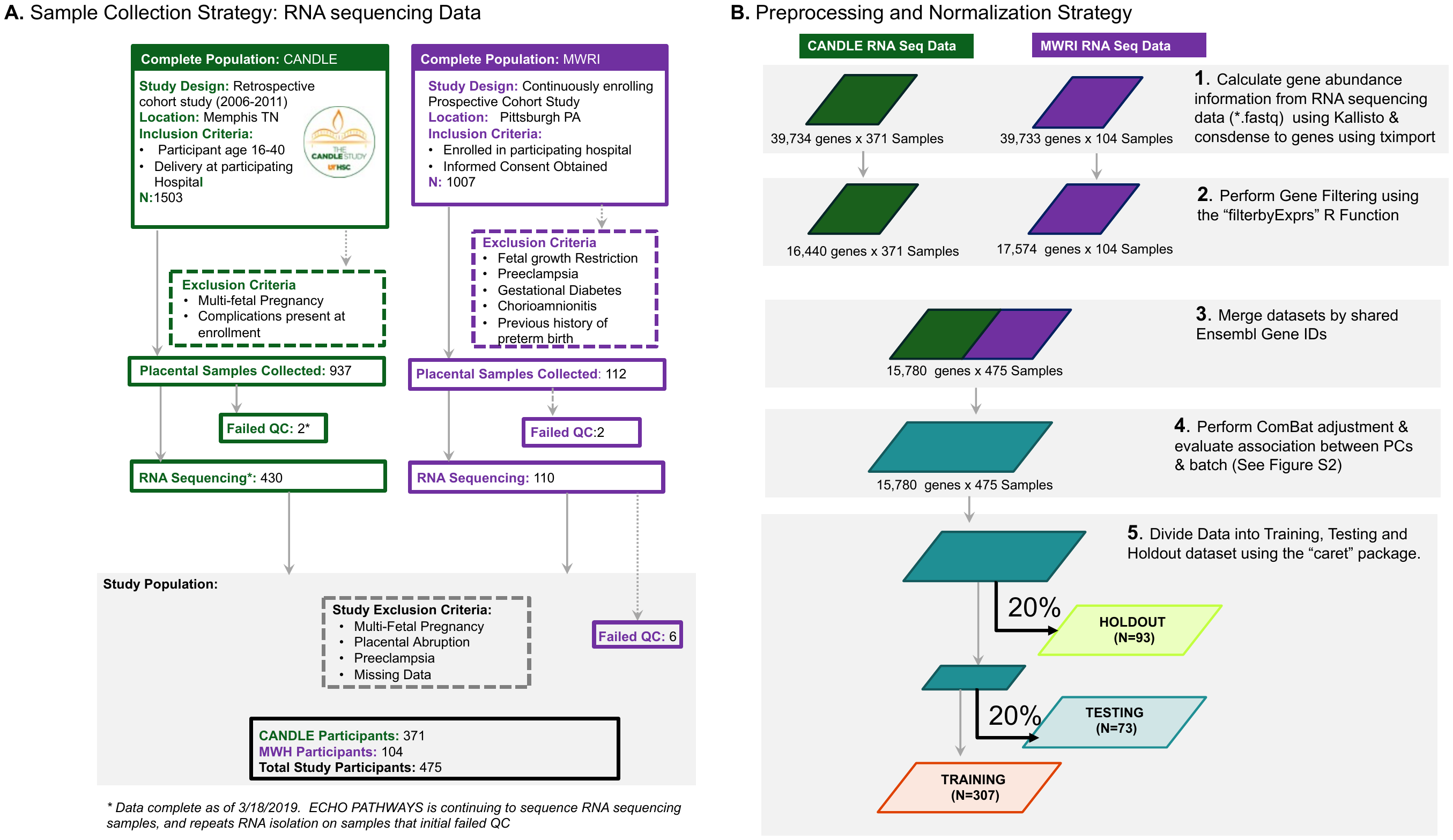

**Supplemental Figure 1**: Flow chart of participant collection (A) and sample processing and dataset creation (B). Samples were recruited from 2 separate populations including the CANDLE cohort, which was a retrospective cohort study taking place from 2006-2011, as well as samples collected from Magee Women’s Research Institute (MWRI) as part of a continuously enrolling study. For the CANDLE samples, we utilized all samples that met study inclusion criteria at the time. We sequenced a subset of samples (112) from all placentas available at MWRI.

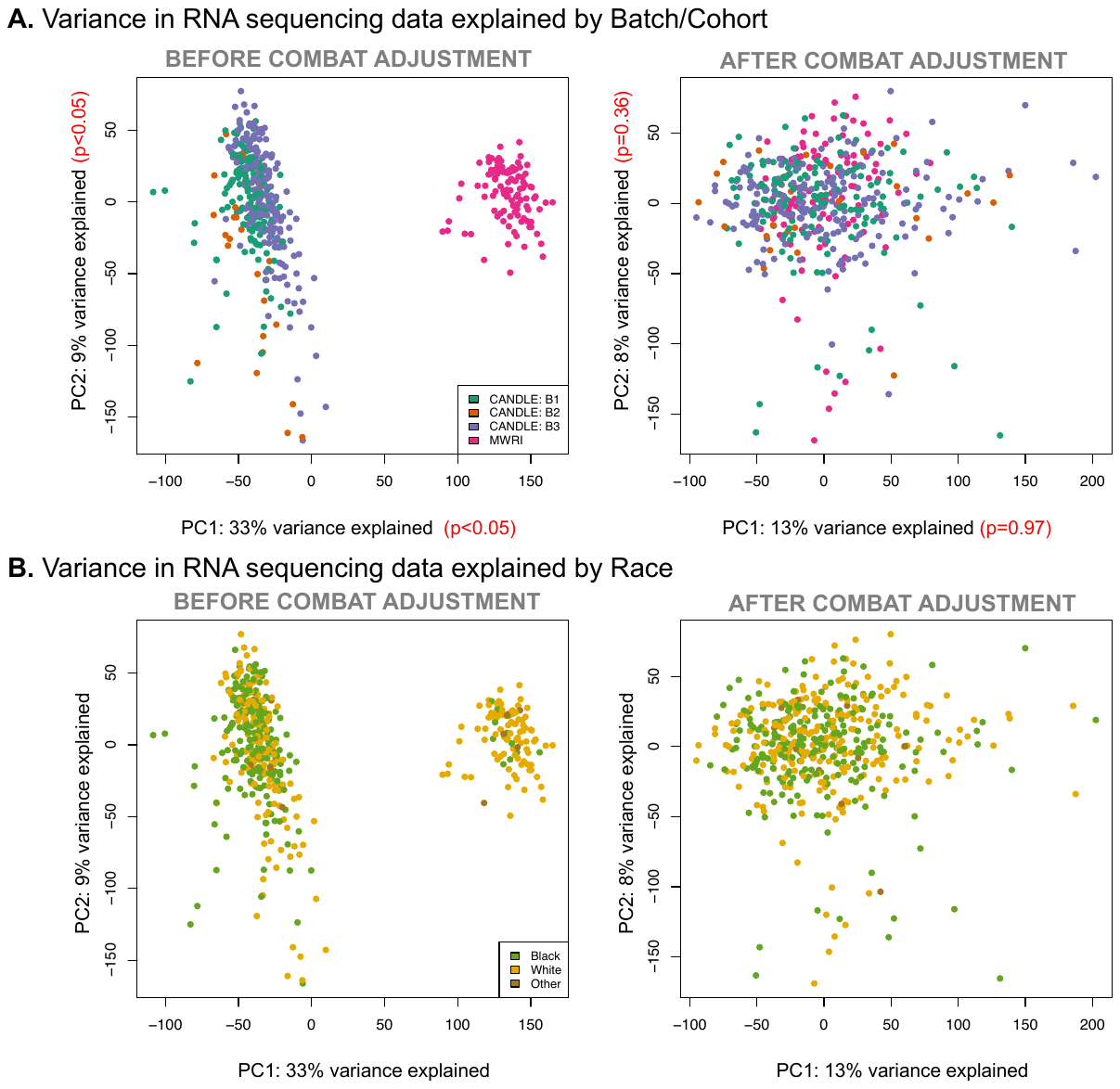

*Supplemental Figure S2:* Relationship between first 2 principal components and RNA sequencing batch/Cohort before and after combat adjustment. P values represent the results of ANOVA on the first and second principal component.

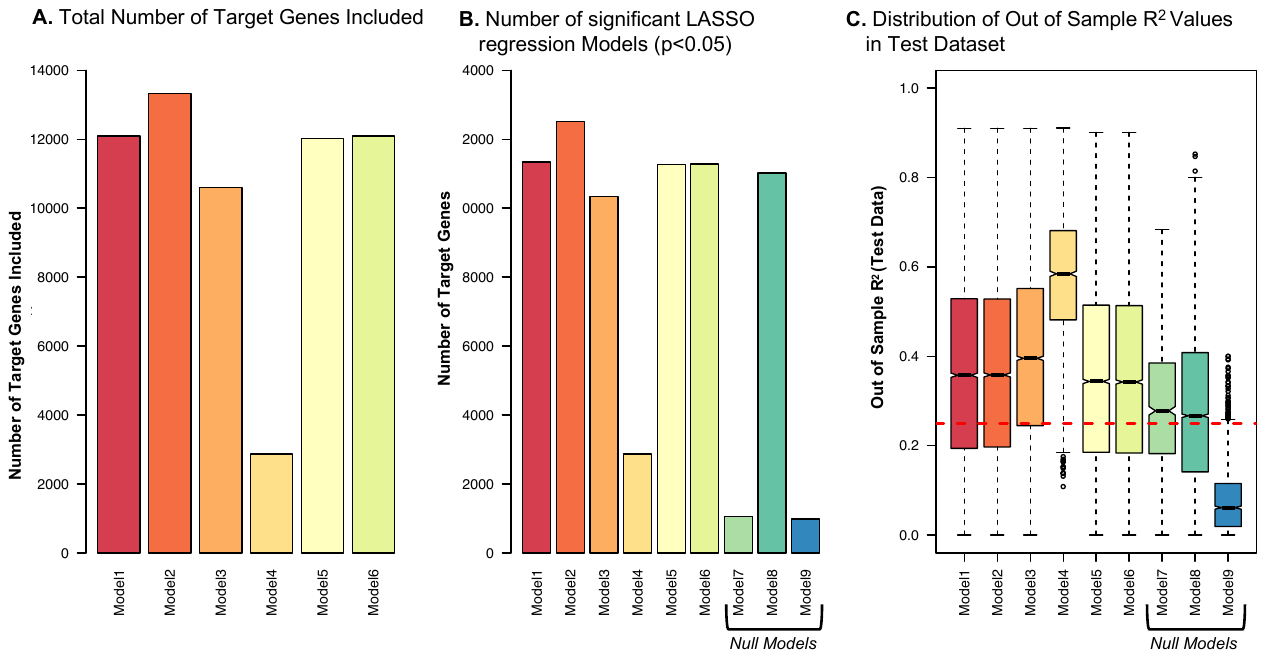

*Supplemental Figure S3*. (A).Number of target genes included in our six primary models after filtering, with full model parameters presented in Supplemental Table S1. (B) Number of target genes with Out of Sample R^2^ values with p> 0.05 in LASSO regression models in test dataset. Here we included 3 null models made with scrambled data-See Supplemental table 2. (C) Range of predictive capacity reflected as out of sample R^2^ value for each gene depicted in model. Out of sample values for 6 permutations of the model using different thresholds for construction and filtering, and 3 null models made of TF-target gene interactions that would not be possible in our actual TRN. Model 1 was used as the reference model. Models 1 and 2 were not statistically different; all other models were (p<0.05, Tukey on ANOVA, Table 4). All null models were significantly different from the true models (p<0.05, Tukey test on ANOVA; See supplemental table 4). Red line represents Out of sample R^2^<0.25, which was used as the cutoff for the final model.

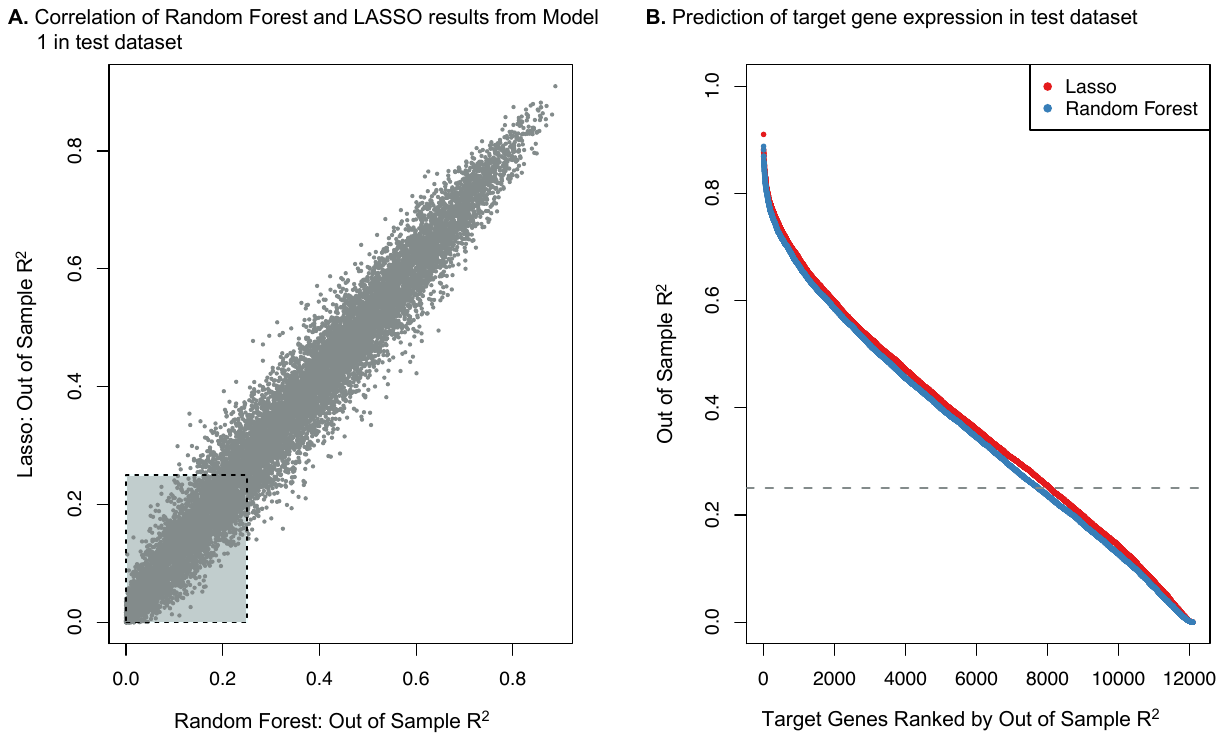

*Supplemental Figure S4.* (A.) correlation between LASSO and random forest OOS R^2^ for 12,094 target genes (from model 1). (Cor=0.97, P <2.2x10-16, Pearson correlation). As shown here, there were few instances where random forest did a substantially better job at predicting gene expression than LASSO regression. The box indicates genes where Random/forest and/or LASSO had a poor prediction accuracy (<0.25) (B.) Target genes (12,368) ranked by OOS R^2^ in both random forest and LASSO regression. Overall; Lasso regression had significantly higher results than random forest (Mean Random Forest R2 =0.38, Mean Lasso R2-0.4,P=1.34x10^-11^, T Test).

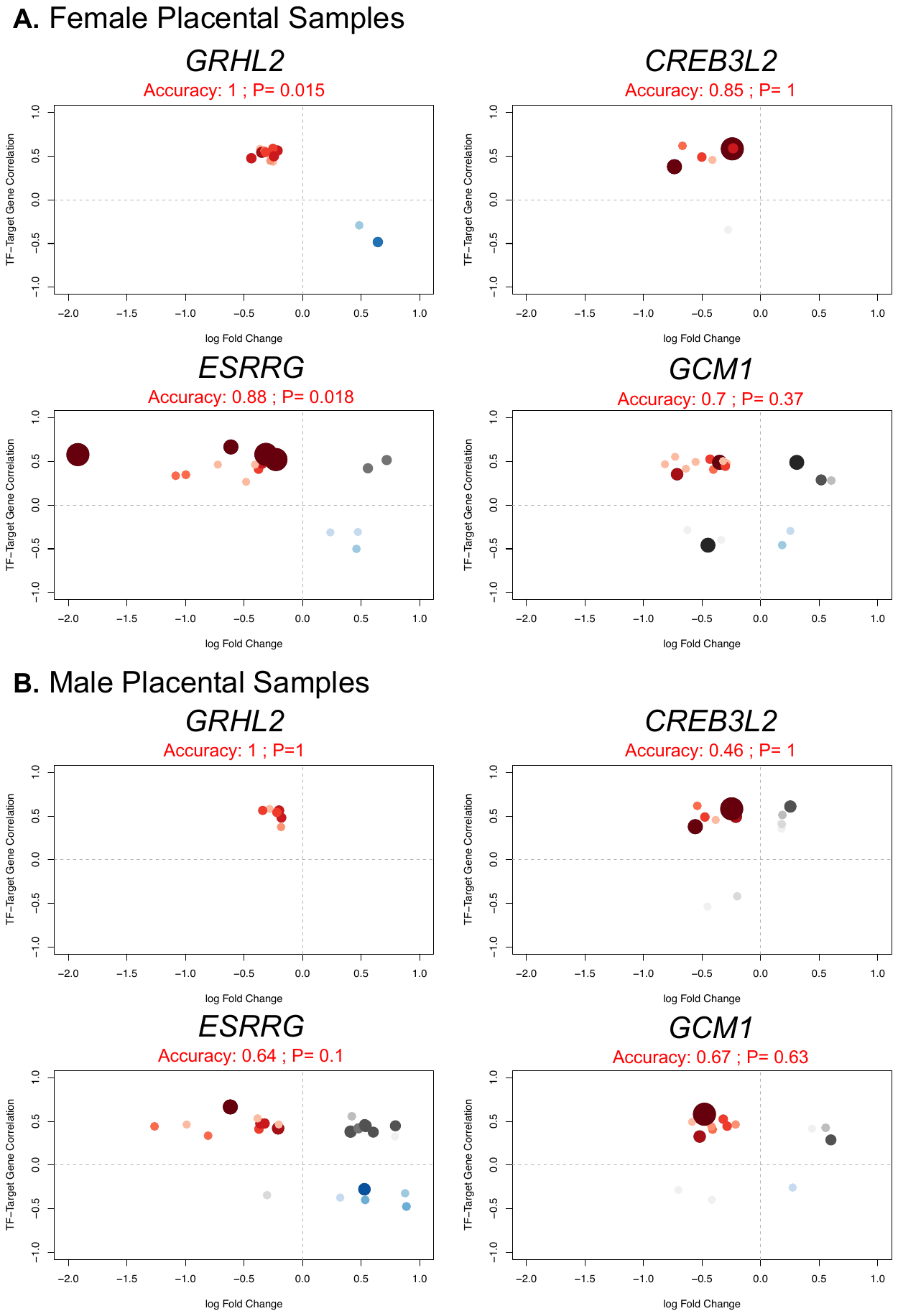

*Supplemental Figure S5:* Results of Experimental validation from figure 4 stratified into (A) female and (B) male samples for the four TFs in our model. The x-axis represents the log fold change of knockout vs control, and, and the y-axis represents the correlation between TF and target gene. Model accuracy was calculated at the number of positively regulated genes with a negative log fold change (i.e “true positives”, shown in red) added to the number of negatively regulated target genes with a positive log fold change (i.e “true negatives”, shown in blue) divided by the total number of target genes for each sample. Genes that were not concordant are shaded in grey. The size and shading of each target gene represents the relative rank (1-15) of the TF as a regulator in our TRN.

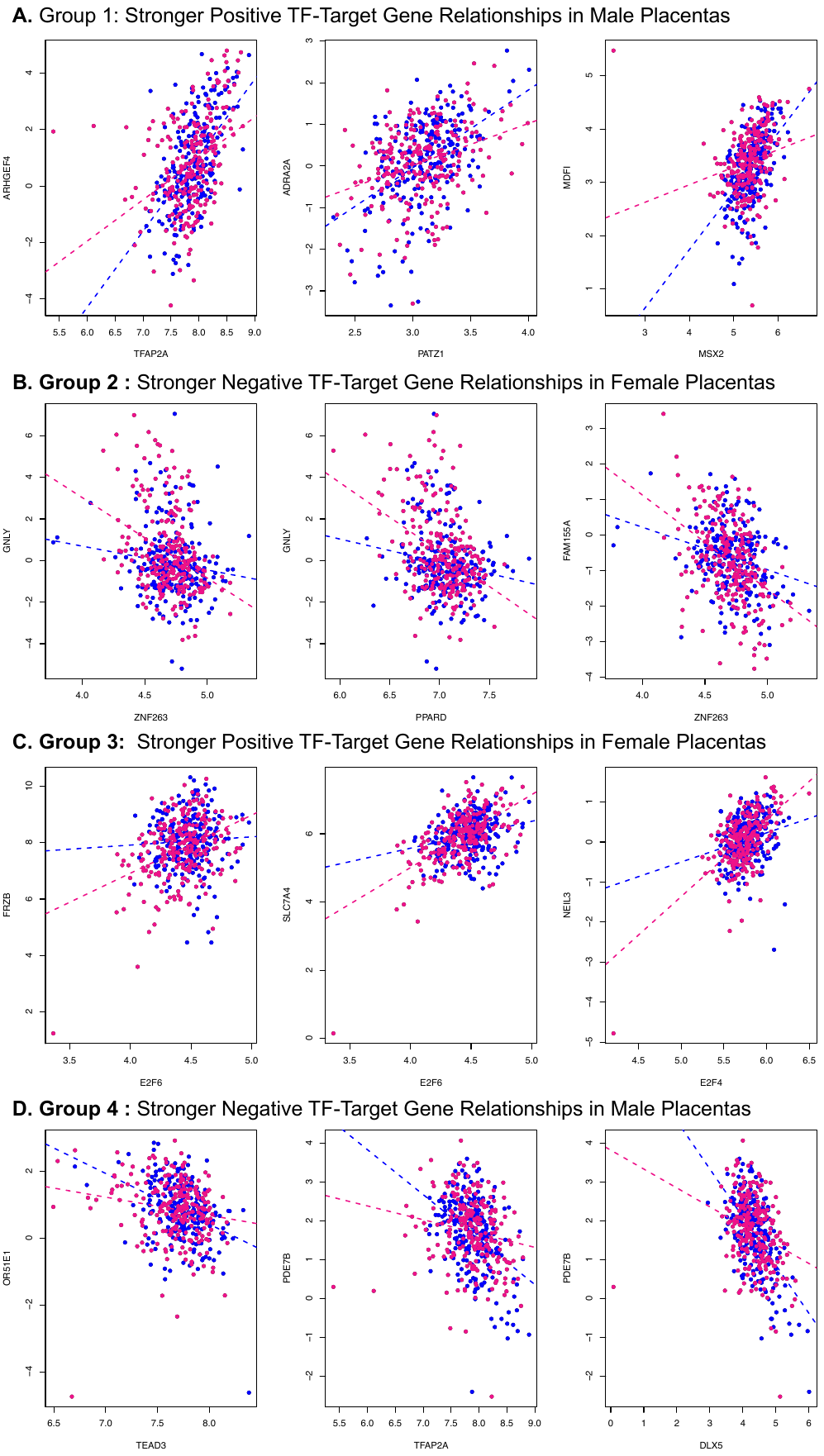

*Supplemental Figure S6:* Top 3 TF-Target gene interactions for each type of TF-target gene relationship including (A.) positive TF-target gene relationships stronger in males than in females (B.) positive TF-target gene relationships stronger in females (C.) negative TF-target gene relationships stronger in females and (D.) positive TF-target gene relationships stronger in males than in females. Gene expression data from females is in pink and in males is blue. Stratified linear models are depicted as line.

**SUPPLEMENTAL TABLE LEGENDS**

*Supplemental Table S1:* Summary of Parameterization variables tested in models constructed in Training and Testing Datasets

*Supplemental* Table S2: Summary of Null Models compared to reference models constructed in Training and Testing Datasets

*Supplemental Table S3:* Results of Tukey test of differences in out of sample R^2^ for each gene in testing dataset.

*Supplemental Table S4:* Summary of regulon (TF) size based on network measures

*Supplemental Table S5:* Table of all significant TF-target gene interaction terms

*Supplemental Table S6:* Full Transcriptional Regulatory Network
